# Radiotherapy drives lung metastasis in PDAC via a RhoGTPase signaling shift towards MRCK-dependency

**DOI:** 10.1101/2025.09.09.673965

**Authors:** KM McLay, M Tesson, L Dutton, Y Sun, JD Misquitta, K Stevenson, R Corbyn, S Lilla, L McGarry, N Maka, R Shaw, C Miller, LM Carlin, AJ Chalmers, MF Olson, JP Morton, JL Birch

## Abstract

Pancreatic ductal adenocarcinoma (PDAC) is an aggressive cancer with an extremely poor prognosis, partly owing to its highly metastatic nature. Although radiotherapy is an effective and potentially curative treatment modality, research over the last few decades has indicated that it may induce a more metastatic phenotype in surviving cancer cells. We demonstrate for the first time that clinically relevant doses of image-guided targeted radiotherapy induce metastasis in a genetically engineered KPC mouse model of PDAC. Furthermore, this induction is largely driven by an organotropic switch towards lung metastasis. Using an *in vitro* RNAi screening approach, we identified a key role for myotonic dystrophy-related Cdc42-binding kinase (MRCK) in driving this response. MRCK activity was spatially upregulated at the plasma membrane in response to radiotherapy in PDAC cell lines as well as at the invasive margins of PDAC tumors. This upregulation of activity was maintained in metastases, suggesting an important role for MRCK in not only triggering local invasion in response to radiotherapy but also promoting distant metastases. Importantly, inhibition of MRCK with a small-molecule inhibitor (BDP9066) specifically opposed radiation-driven MRCK upregulation and pro-metastatic response.

## Introduction

Pancreatic ductal adenocarcinoma (PDAC) is a highly aggressive and metastatic malignancy with a dismal five-year survival rate of less than 10% (1). Despite the low incidence rates, it is projected to become the second leading cause of cancer-related death within the next five years with survival outcomes having not significantly improved since the 1970s, highlighting the urgent need for novel therapeutic strategies (2). PDAC is largely refractory to current treatments, and surgical resection is the only potentially curative option. Chemotherapy offers only modest survival benefits, leading to a dismal median survival of less than six months (3). The role of radiotherapy (RT) in PDAC treatment remains controversial owing to inconsistent clinical trial outcomes regarding its efficacy. While some evidence suggests improved overall survival in the neoadjuvant setting, as demonstrated by the PREOPANC phase III trial, in which a gemcitabine-based chemoradiotherapy regimen resulted in superior survival outcomes compared to upfront surgery and gemcitabine alone in resectable and borderline-resectable cases, other studies have reported conflicting findings (4,5). The LAP07 trial, for instance, demonstrated that although chemoradiation delayed local tumor progression in locally advanced PDAC, it did not confer an overall survival advantage (6). As a result, the current recommendation is for RT to be utilized in patients who are poor surgical candidates or have locally advanced disease (7).

The limited efficacy of RT in PDAC can be partially attributed to its intrinsic radioresistance, which arises from a dense, fibrotic, and immunosuppressive tumor microenvironment with acquired resistance mechanisms that also contribute to treatment failure (8,9). Another factor that may impact the lack of efficacy is RT-induced invasion and metastasis, a phenomenon that has been documented preclinically in multiple malignancies and reviewed in various articles (10–12). However, despite growing evidence supporting this phenotype in other malignancies, studies investigating RT-induced metastasis in PDAC have largely been confined to *in vitro* analyses, with no *in vivo* preclinical studies investigating its impact on the metastatic burden in clinically relevant models (13,14).

RhoGTPases are a family of 26 signaling proteins that drive actin cytoskeletal dynamics to promote cell migration and invasion and consequently play a critical role in tumor metastasis (15,16). Downstream signaling from RhoGTPases is mediated by a panel of effector kinases, including LIM domain kinase (LIMK), Rho-associated protein kinase (ROCK), and myotonic dystrophy kinase-related Cdc42-binding kinase (MRCK), to create a complex and dynamic network (17–19). This complexity leads to a high degree of plasticity that allows cells to adapt to changing conditions within the microenvironment (ME). For example, the RhoA-ROCK signaling axis is used by cells to adopt amoeboid migration to allow navigation through small gaps in the stroma. However, when this form of migration is insufficient to allow cells to pass through dense, complex stroma, cells can switch to a Rac1/CDC42-MRCK driven mesenchymal phenotype that allows extracellular matrix (ECM) degradation and pseudopod formation (20,21).

Interestingly, the RhoA-ROCK axis has been demonstrated to drive invasion in preclinical studies of PDAC under physiological conditions, although how the network responds to radiotherapy to support metastasis remains to be elucidated (19). In glioblastoma, a tumor in which cells adopt a more mesenchymal mode of invasion, MRCK was demonstrated to drive local dissemination of disease under standard of care radiotherapy in preclinical models (22,23). However, the role of other RhoGTPase members and downstream effectors in supporting invasion and metastasis is yet to be comprehensively assessed under physiological or irradiated conditions.

In this study, we demonstrated a pro-migratory phenotype induced by RT in multiple PDAC cell lines. Importantly, we demonstrated for the first time that this increased motility correlates with a significant increase in metastatic burden in a clinically relevant PDAC model, with a tropism switch to the lung. We showed that this effect is mediated by altered RhoGTPase signaling, with RT promoting a dependency on MRCKα signaling to drive the pro-invasive phenotype. Specifically, we showed that RT increases MRCK activity both *in vitro* and *in vivo*, with spatially localized activation at the invasive edge of irradiated KPC tumors and metastases. Furthermore, both RNAi and pharmacological inhibition of MRCK significantly inhibited RT-induced motility, highlighting the potential of anti-invasive strategies for improving RT efficacy. Importantly, given the established role of MRCK in RT-induced invasion of other malignancies, our findings suggest that MRCK signaling may represent a more central mechanism underlying RT-induced invasion across different cancer types.

## Materials and Methods

### Cell culture

cell lines were maintained in incubators at 37°C with 5% CO_2_ , tested regularly for mycoplasma, and authenticated using STR profiling. The PANC-1 (male) and MiaPaCa-1 (female) cell lines were cultured in DMEM (ThermoFisher Scientific #21969035) supplemented with 10% FBS, 1% penicillin/streptomycin (ThermoFisher Scientific 15140-122) and 2mM L-glutamine (ThermoFisher Scientific #25030-024). AsPC-1 cells (female) were cultured in RPMI 1640 medium (ThermoFisher Scientific #11875093) supplemented with 10% FBS, 1% penicillin/streptomycin (ThermoFisher Scientific #15140-122) and 2mM L-glutamine (ThermoFisher Scientific #25030-024). NBC12.2f (female), NBC12.2g (female), and NBC13.3a (male) murine cell lines were generated in-house from C57Bl/6 KPC mice and cultured in DMEM supplemented with 10% FBS, 1% Penicillin/Streptomycin and 2mM L-glutamine. The primary patient lines, TKCC04 and TKCC05, were kindly gifted by Professor David Chang (University of Glasgow), were characterized as the squamous subtype, and were cultured in RPMI 1640 medium (ThermoFisher #11875093) supplemented with 10% FBS, and 20ng/mL Epidermal Growth Factor (Life Technologies #PHG0311L) (24).

### Subconfluent migration assay

Cells (1 × 10^4^ cells/mL) were seeded in an appropriate culture dish for the experimental assay (35 mm dishes or 12-well plates) and left to adhere for 24hours. Drug treatments and irradiation were carried out 4 h prior to imaging unless otherwise stated. Timelapse microscopy images were acquired in phase contrast every 20 mins for 16 hours at 10 X magnification (10x/0.30 Plan Fluor Ph1, Nikon). Cell speed was calculated using single-cell tracking performed in ImageJ. All *in vitro* experiments were conducted in biological triplicates, and statistical analysis details are within the corresponding figure legends.

### Automated cell tracking

Automated tracking was conducted using scripts developed by Dr. Ryan Corbyn (Beatson Advanced Imaging Resource Facility RRID:SCR_023875, CRUK-SI). Time-lapse imaging data were acquired using an IncuCyte S3 system. IncuCyte data was preprocessed to collate the individual time points from the experiment into a single time-lapse image stack followed by drift correction to stabilize the image stacks (Fiji macro “correct 3D drift.”). Bespoke Python scripts were written by Dr. Ryan Corbyn and run in a Jupyter integrated development environment (IDE) for cell segmentation and cell tracking. “We retrained the livecell model from cellpose 2.0 on a subset of the imaging data for automated cell segmentation(25,26). Automated cell tracking was subsequently performed using BTracks, which employs a Bayesian single-cell tracking algorithm and generates deep lineage tree analyses as described by Ulicna et al. ((27)). Tracks of cells present in >90% of the total number of frames were used for further analysis. Further analysis of track lengths and cell speeds was performed for all tracked cells, with the resulting data exported as. csv files.

### High-throughput immunofluorescence

Cells were seeded at a density of 3 × 10³ cells per well in 96-well optically clear plates (Greiner #655892) and allowed to adhere overnight prior to irradiation and/or drug treatments, as specified in the experimental conditions. Treated cells were incubated for an additional 24 h unless otherwise stated. After treatment, the cells were washed with PBS before fixation with 4% formaldehyde at room temperature for 15 min. Cells were permeabilized using 0.1% Triton X-100 in PBS for 10 min, washed with TBS containing 0.1% Tween-20 (TBS-T), and blocked in 1% BSA in TBS-T for 30 min at room temperature. For immunofluorescence staining, cells were incubated with the primary antibody diluted in blocking solution overnight at 4°C. The details of the primary antibodies are listed in Table S1. After primary antibody incubation, cells were washed three times with PBS and then incubated with secondary antibody along with 1:1000 DAPI (Thermo Scientific #D1306), Texas Red-X Phalloidin (Thermo Scientific #T7471) and 1:10,000 HCS CellMask Deep Red Stain (Thermo Scientific #H32721) for 1 h at room temperature. After incubation, the cells were washed and stored in PBS. High-content imaging was performed using the PerkinElmer Opera High-Content Screening System, and image analysis was conducted using the Columbus™ image analysis software.

### Cell viability assays

Cells were seeded at a density of 1 × 10⁴ cells/mL in 96-well plates (Corning) and allowed to adhere overnight. Cells were irradiated at either 0 or 5 Gy using the Xstrahl RS225 and treated with BDP9066 for 24 h. Cell viability was assessed using the CellTiter-Glo® assay (Promega #G9241) and Promega Glomax Multi detection system.

### siRNA transfections

Cells used for transfection were maintained in medium free of penicillin/streptomycin for at least one week prior to the start of experiments. PANC-1 and Mia PaCa-2 cells were transfected with siRNA targeting CDC42BPA (MRCKα) or control scrambled siRNA (Dharmacon #L003290-00; MRCKα AAGAAUAUCUGCUGUGUUU) using DharmaFECT 2 transfection reagent (Dharmacon #T-2005-01) and incubated for 48 h prior to imaging or protein extraction.

### Proteome analysis

Fifteen-centimetre dishes were pre-coated with Matrigel prior to cell seeding to increase cell adherence. Matrigel was diluted at a ratio of 1:40 in cold DMEM medium without additional supplements. A total of 5 mL Matrigel dilution was added to each dish and incubated at 37°C and 5% CO₂ for 1 hour. Excess medium was aspirated prior to cell seeding.

PANC-1 cells were seeded at a density of 2 × 10⁵ cells/mL in Matrigel-coated dishes, with 15 mL of cell suspension per dish. Three biological replicates were plated. Cells were left to adhere overnight at 37°C and 5% CO₂, resulting in ∼80% confluency. Six hours before analysis, the media were aspirated, and plates were washed six times with PBS to ensure all cell debris was removed. Media was then replaced with 10mL FBS-free culture medium. Immediately after media replacement, cells were irradiated with either 0 Gy or 5 Gy and incubated for 6 hours. Following this incubation, cell lysates were collected for proteome and secretome analysis, respectively.

Ultra-High Performance Liquid Chromatography/Tandem Mass Spectrometry (UHPLC-MS/MS) analysis and data analysis conducted by the CRUKSI proteomics facility. See supplemental data for details.

### RNAseq

NBC12.2g, NC12.2f and NBC13.3a were irradiated +/-5 Gy. RNA was extracted 24 hours post irradiation using Rneasy kit (Qiagen) kit. Truseq (Illumina) mRNA-seq libraries were prepared from total RNA and these were then sequenced using the Illumina HiSeq 4000 using 50M read pairs, 2×75bp read length. DEseq2 was utilised for differential gene analysis. Volcano plot created in GraphPad Prism v10.60. See supplementary methods for full analysis details.

### RNAi screen

PANC1 cells were seeded at a density of 1×10⁵ cells/mL in two 12-well plates and allowed to adhere overnight. Cells were transfected with 25 nM custom siRNA ON-TARGETplus SMARTpool library (Horizon) using DharmaFECT 2 transfection reagent and incubated overnight. The details of the siRNA are shown in Table S2. After incubation, transfected cells were washed twice with PBS, detached from the wells using TrypLE (250 µL per well), and resuspended in an additional 750 µL of antibiotic-free medium. Cells were plated at a density of 1×10⁴ cells/mL in 24-well plates for the subconfluent migration assay. In parallel, selected transfected cells were replated into additional 6-well plates for protein extraction to verify successful knockdown. All replated cells were allowed to adhere overnight at 37°C and 5% CO₂ and subject to 0 Gy or 5 Gy after 16hrs and 4 h prior to imaging.

### Western blot

Cells were lysed with 1% SDS and 50 mmol/L Tris (pH 6.8) supplemented with protease and phosphatase inhibitors (Thermo Scientific #1861280) followed by SDS-PAGE using 4–12% Novex Bis-Tris gels or 8% BOLT Bis-Tris Plus gels (Thermo Fisher Scientific), and transfer onto nitrocellulose membranes and western blotting with enhanced chemiluminescence (ECL) detection. Primary antibodies were diluted in 5% bovine serum albumin (BSA). The details of the primary antibodies used are shown in Table S1.

### *In vivo* experiments

All animal work was conducted under the guidance of the UK Home Office regulations and carried out under project license PP8411096. All experiments were approved by the University of Glasgow Animal Welfare and Ethical Review Board. The mice were housed in standard cages with access to food and water *ad libitum*. Environmental enrichment was also provided. KPC mice of both sexes were used in all the experiments. *Pdx1-Cre*; *LSL-Kras^G12D/+^*; *LSL-Trp53^R^*^172^*^H/+^* (KPC; (28)) mice were bred in-house at the CRUK Scotland Institute, maintained on a mixed background, and genotyped using Transnetyx (Cordoba, TN, USA). The mice were palpated weekly to detect pancreatic tumors, and the presence of a pancreatic tumor was confirmed by 3-dimensional (3D) ultrasound using the Vevo3100 high-frequency ultrasound platform (FUJIFILM VisualSonics) prior to enrolment in the treatment arm. Mice in the treatment group were housed in the same room, and single housing was avoided where possible. In the gemcitabine study, mice were intraperitoneally administered 100 mg/kg gemcitabine twice a week.

### *In vivo* irradiation of pancreas tumour

The Small Animal Radiotherapy Research Platform (SARRP, Xstrahl) was used to deliver targeted irradiation as previously described (29). Mice were anesthetized using a mixture of medical air and isoflurane for the duration of the radiotherapy session. An iodine contrast-enhanced cone beam computed tomography (CBCT) image was obtained from 1,440 projections using the in-built cone beam CT function of SARRP. To enhance contrast, the mice were administered 0.25 ml Iohexol (Omnipaque™, GE Healthcare) intraperitoneally immediately prior to CT acquisition. Radiotherapy planning was performed using Muriplan software, including tissue segmentation and placement of an isocenter to localize the pancreatic tumor. Arc radiotherapy was delivered at 280 cGy/min (220 kV, 13 mA) using three fractions of 4 Gy every other day, three fractions of 6 Gy every other day, or a single fraction of 12 Gy. The collimator aperture size was selected based on tumor shape and size. The mock irradiation controls consisted of mice subjected to 1,440 CBCT acquisitions, which resulted in the delivery of 6.22 cGy (29). Mice were monitored by researchers blinded to the treatment arm and euthanized when they exhibited clinical signs of PDAC (distended abdomen, loss of body conditioning, lethargy, and jaundice).

### Histology and immunohistochemistry

Tissue samples were fixed in formalin and embedded in paraffin using standard protocols before acquisition of 4 μm sections using a HistoCore MULTICUT microtome (Leica, UK). Tissue sections were placed on poly-L -lysine slides and oven baked at 60 °C for 2 h prior to staining with hematoxylin and eosin (H&E) or immunohistochemistry (IHC) for Zeb1, and pMRCK. The full list of antibodies used is presented in Table S1.

All liver and lung tissues from control mice and those that received 3 × 4 Gy of radiation were subjected to serial sectioning (five sections at 50 µm intervals). The sections were blinded and double scored for the presence of metastases. For IHC, antigen retrieval and antibody dilution were optimized in-house for each antibody (Table 2.8).

Automatic quantification of staining was performed using QuPath ((30), v0.05.1). The full slides were scanned and imaged using a Lecia Aperio AT2 slide scanner. Quantification was conducted blinded, with either the full tumor or five areas at the tumor center and tumor border selected at random. These areas were annotated using QuPath annotation tools. The percentage of positive cells was then calculated.

### Statistical analysis

Statistical analyses are detailed in the figure legends for all experiments. Statistical analyses were performed using the GraphPad Prism software. The number of independent experiments for each in vivo experiment is within the figure legend and refers to experiments repeated on different days with cells at different passages for each cell line reported. Biological repeats refer to independently derived cell lines used as replicates.

## Results

### Radiotherapy induces motility across a panel of PDAC cell lines

Initial qualitative morphological analyses indicated that sublethal doses of RT (5 Gy) induced a shift in morphology of PANC1 cells towards a more elongated shape, reminiscent of a mesenchymal transition (Fig. 1A). Indeed, comprehensive and unbiased high-throughput immunofluorescent imaging and automated analysis indicated that radiotherapy significantly increased the cell area in two PDAC cell lines, which may be indicative of a motile phenotype through increased engagement with integrin signaling (Fig. 1B)(31). Previous studies have reported increased PDAC cell motility *in vitro* in response to RT; however, these studies have been limited in terms of cell line number and available technologies, and as such, have yielded conflicting results ((32–34); as summarized in Table S3). Therefore, we used a comprehensive approach across a panel of commercial, mouse-derived, and primary patient cell lines to measure the effect of RT on PDAC cell motility. We used clinically relevant doses of RT (2 and 5 Gy) combined with subconfluent migration assays and single cell tracking to eliminate potential confounding effects on proliferation. We identified a significant increase in migration speed across the selection of three commercial, mouse, and primary lines (Fig. 1C). Parallel cell viability assays indicated that RT had a modest impact on the viability of the TKCC04 line only at this short timepoint (24hrs; Fig.1D). Expansion of this panel indicated that, while the effect of RT on migration is not ubiquitous, five out of eight lines showed increased migration speed post RT, with a further two showing a trend towards an increase (Table 1). Of note, the only murine line that did not show a significant increase in speed, NBC12.2g, was more radiosensitive than the other two lines (Fig. S1).

**Fig 1.**
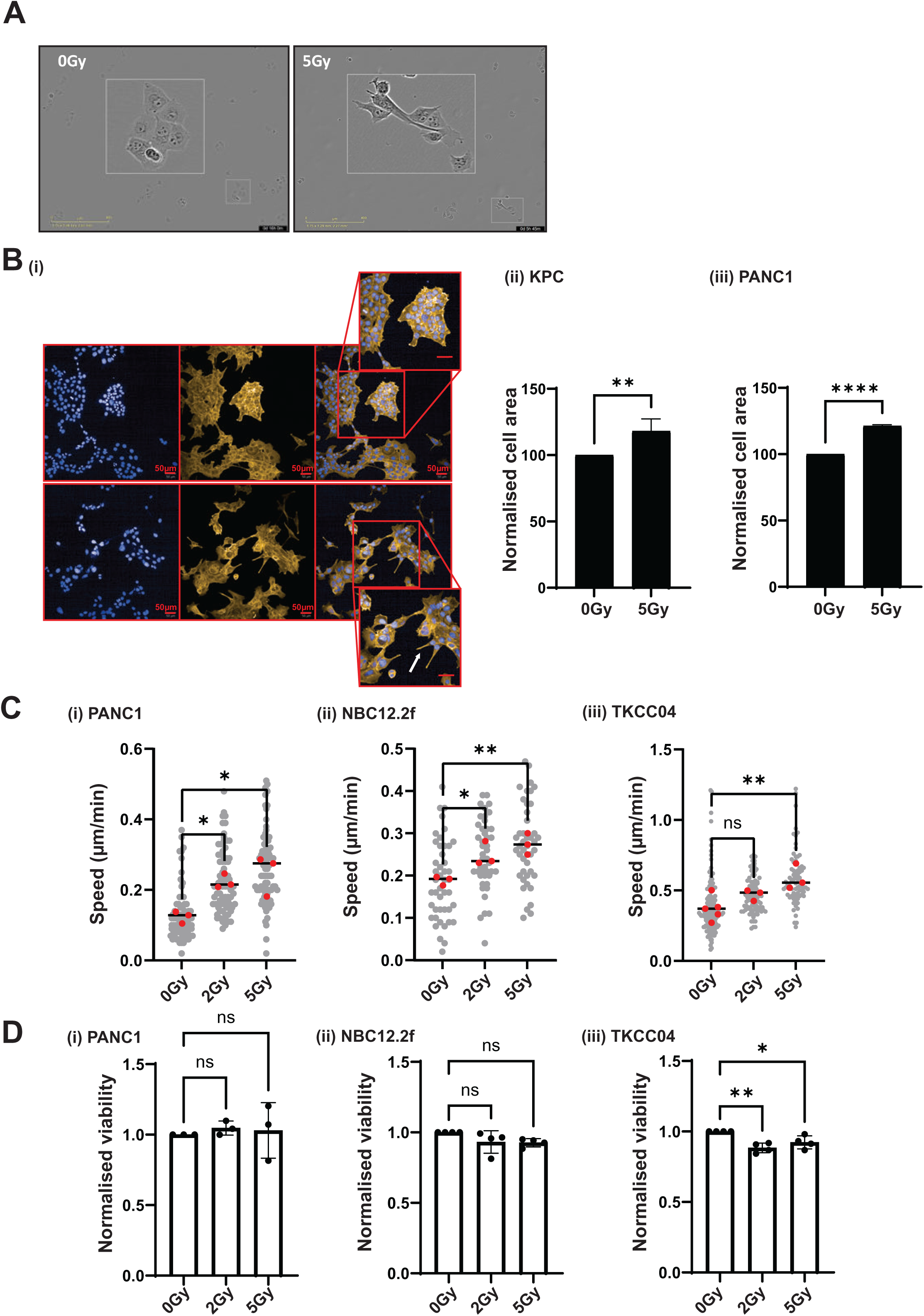
Radiotherapy induces cell motility across a panel of PDAC cell lines. **A)** PANC1 cells imaged under phase contrast indicating morphology shift following exposure to 5 Gy RT. **B) (i)** PDAC cells treated with either 0 Gy or 5 Gy RT and imaged via high-throughput imaging. (Blue, DAPI; yellow, actin; last panel, merge) followed by automated analysis of cell area **(ii)** KPC (NBC12.2f) **(iii)** PANC1. Scale bar, 50 μm. Data derived from three independent experiments each containing 4 internal replicates, >1000 cells measured per condition per independent experiment. Error bars = Standard Deviation. Statistical analysis: two-tailed, unpaired *t* test. **, *P <* 0.01; ****, *P* < 0.0001. **C) (i)** PANC1 **(ii)** NBC12.2f (KPC) and **(iii)** TKCC04 cells treated with 0, 2 or 5 Gy RT followed by subconfluent migration assay 24 hrs post RT using time-lapse microscopy and single cell tracking to quantify migration speed. Individual cell speeds plotted in grey, means from independent experiments plotted in red, mean from independent repeats = horizontal bar. Statistical analysis: One-way ANOVA; ns, not significant, *, *P* < 0.05, ** *P* < 0.01. **D)** Viability assays conducted 24hrs post RT on **(i)** PANC1 **(ii)** NBC12.2f and **(iii)** TKCC04. Data from three independent experiments for each cell line. Statistical analysis: One-way ANOVA; ns, not significant, *, *P* < 0.05, ** *P* < 0.01.

**Table 1:** Migration speed responses to 2 or 5 Gy in a panel of PDAC cell lines. The migration speed was normalized to 0 Gy to determine the fractional change. Statistical analysis: Student’s t-test. Green cells: significant increase in speed post RT, blue cells: trend towards increase, white cells: no increase. Data were obtained from three biological replicates for each cell line.

| Cell line | Origin | 2Gy |  | 5Gy |  |
| --- | --- | --- | --- | --- | --- |
| | | $\Delta$ speed | P value | $\Delta$ speed | P value |
| PANC1 | Commercial | 1.822849 | 0.0027 | 1.98868 | 0.0024 |
| Miapaca2 | Commercial | 1.07396 | 0.6494 | 1.019541 | 0.8727 |
| ASPC1 | Commercial | 0.879926 | 0.3841 | 1.267169 | 0.0658 |
| NBC12.2f | Murine (KPC) | 1.317871 | 0.0163 | 1.451858 | 0.0002 |
| NBC12.2g | Murine (KPC) | 1.327689 | 0.0618 | 1.044459 | 0.7157 |
| NBC13.3a | Murine (KPC) | 1.692982 | <0.0001 | 2.056329 | 0.0049 |
| TKC004 | Primary patient derived | 1.455445 | 0.0085 | 1.413566 | <0.0001 |
| TKC005 | Primary patient derived | 1.58574 | 0.0137 | 1.558938 | 0.0099 |

### Radiotherapy increases the metastatic burden of pancreatic cancer in a clinically relevant genetic mouse model

While the activation of cancer cell migration is the initiating step in metastasis, a number of key processes are required within the context of an entire organism, including local extracellular matrix invasion, dissemination via the vasculature/lymph system, and seeding of distant sites. In light of this, we assessed whether the increased motile phenotype translates to an increased metastatic burden *in vivo* in a genetically engineered KPC mouse model of PDAC (*LSL-Kras*^G12D/+^; *LSL-Trp53*^R172H/+^; *Pdx1-Cre*) in combination with image-guided targeted RT using a Small Animal Irradiation Platform (SARRP) (Fig. S2A). This model simulates the entire metastatic cascade following targeted irradiation of the primary tumor, allowing us to assess the response to RT in a clinically relevant model and to gain insight into the potential clinical impact of RT on metastasis (Fig. 2A). To ensure a robust analysis, we cut five H&E sections of liver and lungs from control and irradiated (3 × 4 Gy) PDAC mice at 50µm intervals, which were blinded and double scored for the presence of metastases (Fig. 2B). The results indicated that RT induced a significant increase in not only local invasion (Fig. 2C) but also metastatic burden (Fig. 2D). Importantly, 3 × 4 Gy did not prolong the survival of KPC mice, as shown in Fig. S2B and previously published data (29), ruling out the possibility that the radiation-associated increase in metastasis may be due to the additional time for tumor cells to seed metastases. In agreement, there was no association between the lifespan of the animal and the presence of metastases on post-mortem examination (Chi Square, *P*=0.564).

**Fig 2.**
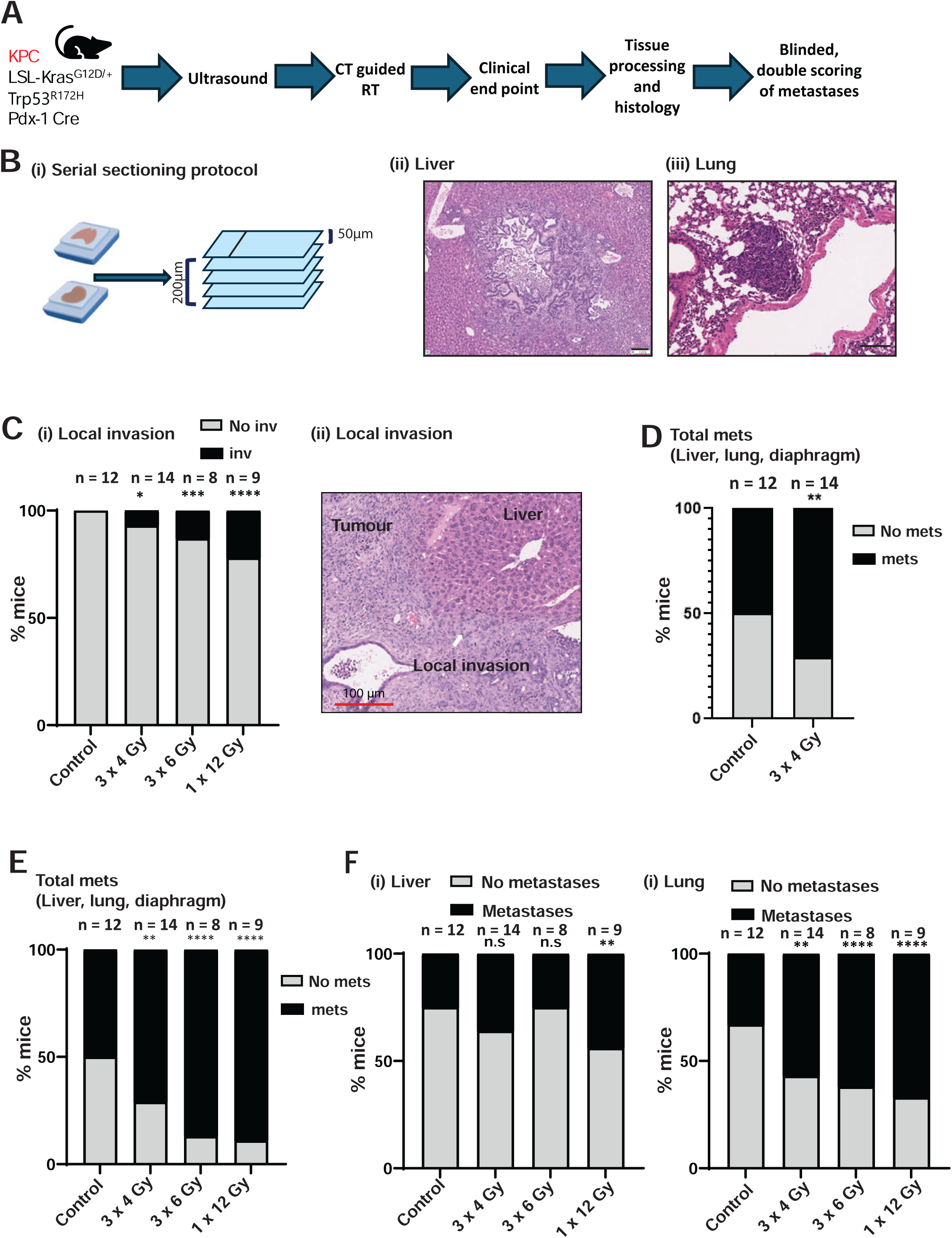
Radiotherapy increases the metastatic burden of pancreatic cancer in a clinically relevant genetic mouse model. **A)** Tumour bearing KPC mice were subjected to CT-guided targeted RT and sampled at clinical endpoint. Liver and lung sections were blinded and double scored for presence of metastases. **B) (i)** Serial sectioning protocol **(ii)**, **(iii)** example images of liver and lung metastases. **C)** i) H&E sections from KPC mice treated with 0Gy, 3×4Gy, 3×6Gy or 1×12Gy were scored for the presence of local invasion (example in (ii)). **D)** H&E liver, lung, and diaphragm sections from KPC mice treated with 0 or 3×4Gy RT were assessed for presence of metastases. **E)** Metastasis scoring across multiple RT protocols, as indicated. **F)** Breakdown of metastases present in liver **(i)** or lung **(ii)**. Number per cohort indicated on graphs. Statistical analysis: Chi-square test. ns, non-significant, *, *P* < 0.05, **, *P* < 0.01, ***, P< 0.001 ****, *P* < 0.0001.

Radiotherapy is given in fractionated doses in the clinic, with a trend towards hypofractionation in recent years. To assess whether fraction size and number impact the effect of RT on metastasis, we extended our analysis to include cohorts of mice that received 3 × 6 Gy (increased fraction dose) and 1 × 12 Gy (hypofractionation). However, only a significant increase in survival was observed in the 1 x 12 Gy cohort (Fig. S2B), all three RT protocols showed a significant pro-metastatic outcome (Fig. 2E), whereas mice treated with gemcitabine alone showed a reduction in metastatic burden (Fig. S2C). These data suggest that RT promotes metastasis independent of the relative lifespan of the treated animals.

PDAC metastasizes primarily to the liver and lungs in both patients and the KPC model. Interestingly, and in support of a radiation-specific metastatic induction mechanism, when we analyzed lung and liver metastases separately, we found that RT induced an organotropic switch towards the lungs across all irradiated cohorts (Fig. 2F).

### RT augments RhoGTPase signaling towards a dependence on MRCK and RhoU

To unpick the potential mechanisms that drive the rapid motility response that we observed post RT and begin to understand how this might trigger the metastatic cascade, we performed RNA-seq and proteomics on PDAC lines post RT/sham RT. There were relatively few genes (at 24h post RT) or proteins (6h post RT) that displayed significant fold changes greater than 1.5 at the relatively short time points used, and no evidence of an emerging motility signature (Fig. 3A-B). This was the case when the cut-off was reduced to a 1.2-fold change in the proteomic data, with response to cellular stress emerging as the main signature post RT (data not shown). Therefore, we sought to undertake a more targeted functional screen of known drivers of motility that are regulated rapidly and independently of gene expression. RhoGTPases drive cancer cell motility and invasion through direct action on the cytoskeleton and are therefore likely to be required to support the observed changes in motility and metastasis.

**Fig 3.**
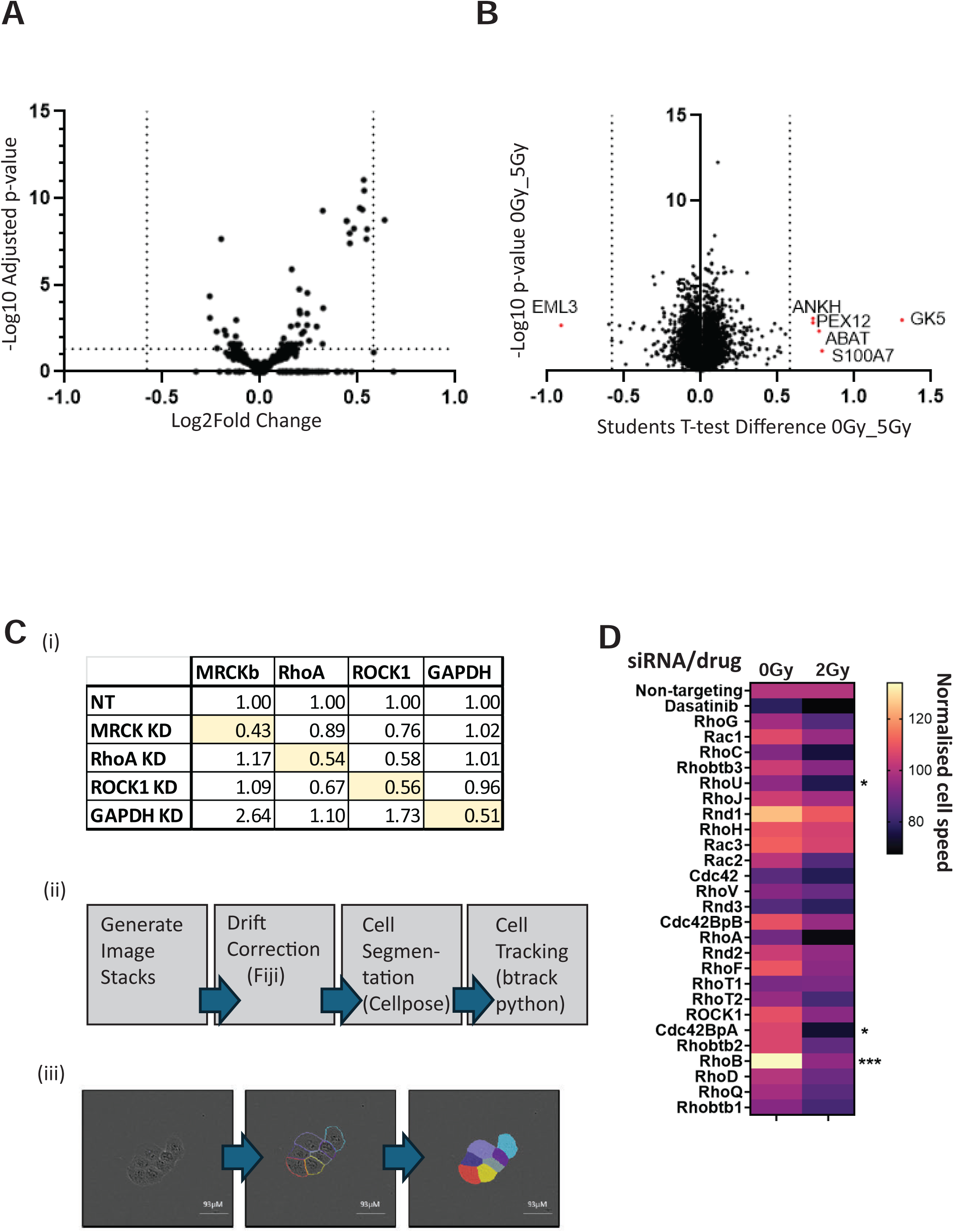
Dependence of RT induced motility on members of the RhoGTPase signaling family. **A-B)** Differential gene expression (DGE) was analysed 24 hrs post treatment with 0 or 5Gy RT in three biological replicates (independently derived KPC lines, NBC12.2f, NBC12.2g and NBC13.3a). The proteome of PANC1 cells was analysed using a tandem mass tag approach 6 hrs post treatment with 0 or 5Gy RT. **C (i)** Efficacy of knockdown was confirmed by western blotting for several targets. Expression normalized to actin loading control and shown as ratio of non-targeted control treated cells (NT). Example blot in Fig. S3. **(ii)** Automated cell tracker workflow. Generation of image stacks and application of drift correction via plugin in Fiji followed by cell segmentation using cellpose 2.0 and tracks cells using btrack python package. **(iii)** representative image of cell segmentation from cellpose 2.0. **D)** RhoGTPase siRNA motility screen targeting all members of the RhoGTPase signaling family and select effector kinases in PANC1 cells exposed to either 0 or 5 Gy RT. Presented as a heatmap where each cell speed is normalised to non-targeting control siRNA from the match condition (0 or 5Gy). Statistical analysis: Two-way ANOVA. *, *P* < 0.05, ***, *P* < 0.001. Statistical analysis: two-tailed, unpaired *t* test. ns, non-significant, *, *P* < 0.05, **, *P <* 0.01; ****, *P* < 0.0001.

We developed an unbiased, AI-driven, automated segmentation and single-cell tracking approach to assay the dependence of PDAC cells on RhoGTPase family members and associated kinases pre-and post RT for migration (Fig. 3C). To mitigate the inherent plasticity and redundancy in the RhoGTPase network, we used an siRNA approach to transiently knock down each target in PANC1 cells. Using this approach, we achieved knockdown efficiency of approximately 50% across a range of selected targets (Western blot analysis; Fig. 3C(i); Fig. S3). The results in Fig. 3D indicate that there is an overall shift towards dependency upon RhoGTPase activity for motility following RT, with a significant interaction between the presence or absence of RT and the ablation of RhoU, RhoB, and MRCKα (CDC42BPA) expression. Closer examination of the data revealed that while the RhoB interaction was driven by dependency under control conditions, PANC1 cells showed a shift in dependency towards RhoU and MRCKα following RT, suggesting that they may play a role in driving RT-induced motility (Fig. 4A). We used a second siRNA oligo sequence coupled with manually tracked subconfluent migration assays to validate the dependency of irradiated PANC1 cells on MRCKα for motility (Fig. 4B(i), (ii)).

**Fig 4.**
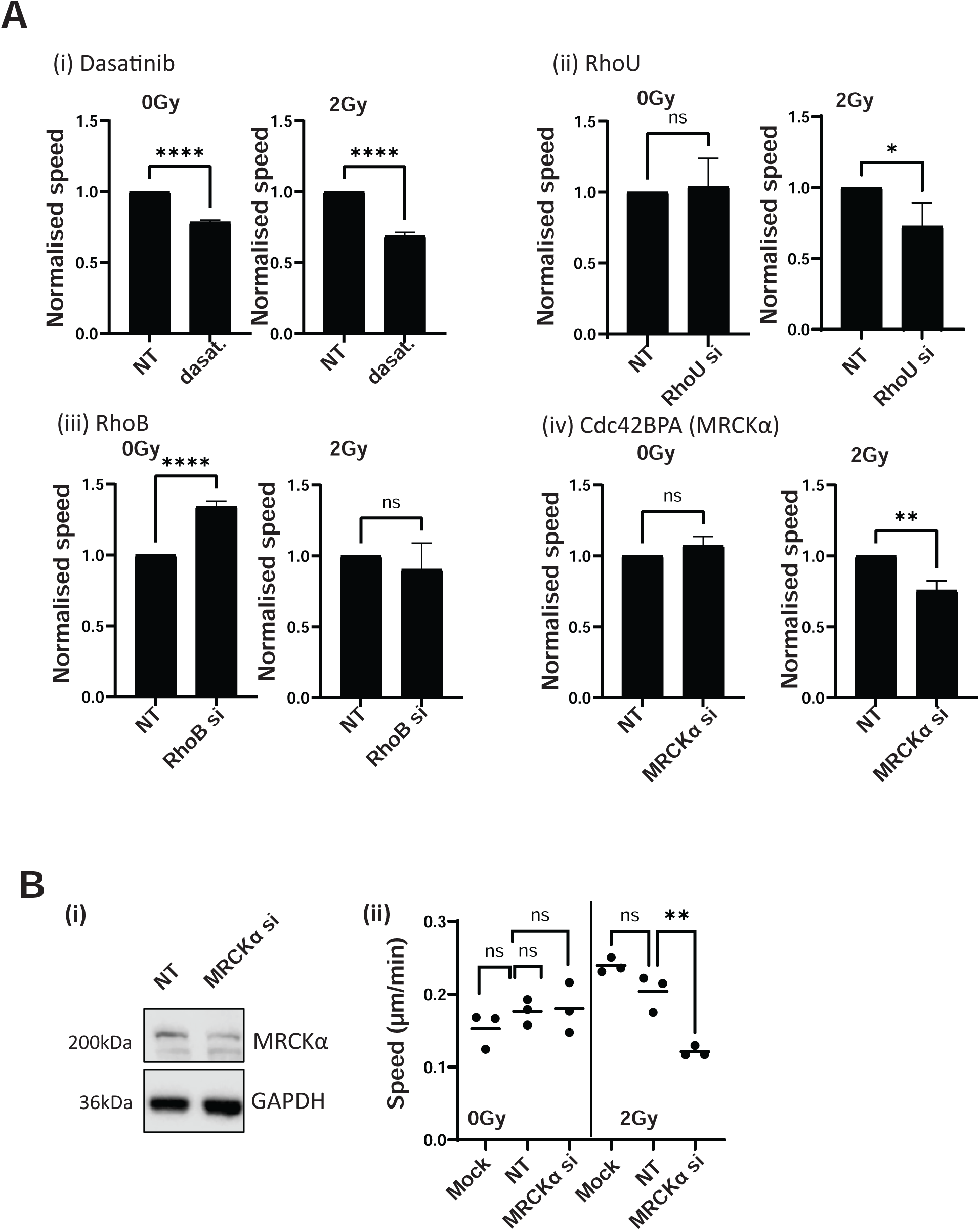
RT-induced motility is dependent on MRCK and RhoU. **A)** Normalised cell speeds of identified hits from the RhoGTPase siRNA screen. **(i)** Dasatinib (positive control) **(ii)** RhoU **(iii)** RhoB and **(iv)** Cdc42BPA (MRCKα). All data from 3 independent experiments with >50 cells tracked per condition per experiment. **B)** PANC1 cells transfected with an alternate siRNA targeting MRCKα followed by **(i)** Confirmation of MRCKα knockdown via western blotting and subconfluent migration assay. N=3 independent repeats, horizontal bar = mean. Statistical analysis: one-way ANOVA. ns, non-significant, **, *P* < 0.01.

### The induction of spatially regulated MRCK signaling by RT is conserved *in vitro*

Metastasis is a leading cause of morbidity and mortality in pancreatic cancer and represents an urgent and unmet clinical need. Given the availability of small-molecule inhibitors targeting MRCK, we selected this kinase for further validation (35). Analysis of available patient datasets also indicated that higher MRCKα mRNA expression showed a trend towards association with poorer overall survival (p = 0.059; Fig. 5A(i)), unlike MRCKβ, which has no association with survival. MRCK and related kinases are well described to be regulated by phosphorylation and intracellular location, which may explain the weak interaction between outcome and RNA expression in patient datasets. In support of this, preliminary single-cell RNA-seq data from KPC mice indicated a significant but minor increase in MRCKα expression post RT (data not shown).

**Fig. 5.**
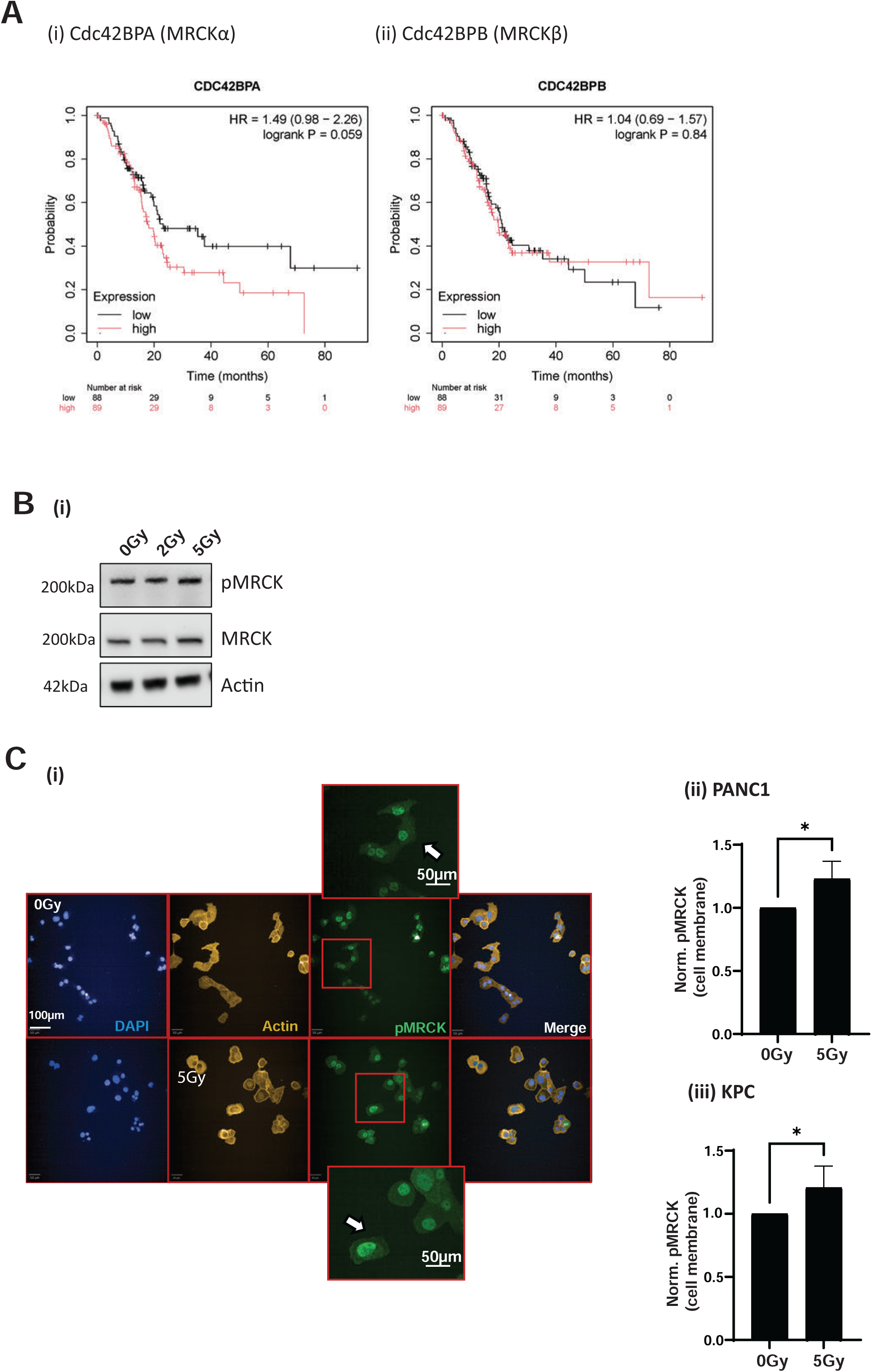
MRCK signaling is spatially regulated by RT *in vitro*. **A)** CDC42BPA (i) and CDC42BPB (ii) mRNA expression and overall survival from available patient datasets plotted using kmplot.com. 177 samples with a median expression cut off. **B)** PANC1 cells treated with 0, 2Gy or 5Gy followed by western blotting of pMRCK total MRCK and actin (loading control). **C) (i)** PANC1 cells treated with either 0 Gy or 5 Gy RT followed by IF for pMRCK and high throughput confocal imaging. Blue, DAPI; yellow, actin; green, pMRCK; last panel, merge. Scale bar, 100 μm. (ii) automated image analysis of membrane associated pMRCK in **(ii)** PANC1 **(iii)** KPC (NBC12.2f) cells exposed to 0 or 2 Gy. All data from 3 independent repeats. Statistical analysis, Students T Test. * *P* < 0.05.

To explore this, we performed a western blot analysis to validate the biomarker of MRCK activity (autophosphorylation of S1003; pMRCK; Fig. 5B). Using the PANC1 cell line, a highly motile cell line in response to radiation (Fig 1C), we found no increase in pMRCK levels in response to 2 or 5 Gy RT (Fig 5C), suggesting that spatial regulation of RhoGTPase signaling may be a more useful determinant of downstream signaling and the resultant phenotype. Therefore, we utilized high-throughput immunofluorescence imaging with automated analysis to examine MRCK activity within different cellular compartments. We demonstrated that MRCK activity significantly increased at the cell membrane following irradiation across the two cell lines, indicating the induction of discrete cellular responses to promote migration (Fig. 5C).

### Pharmacological targeting of MRCK with a small molecule inhibitor opposes radiation driven migration

We used a potent and specific small-molecule inhibitor of MRCK, BDP9066, to test the potential therapeutic benefit of targeting MRCK in combination with RT in PDAC (35). The results in Fig. 6A (i) –(iii) indicate that treatment of PANC1 cells with BDP9066 significantly reduced pMRCK at the cell membrane only under irradiated conditions, demonstrating a shift towards MRCK dependency post RT. ROCK inhibitor (Y27632) was used as a negative control. Furthermore, the data in Fig. 6A (iv) revealed a dose-dependent reduction in pMRCK in response to BDP9066 with a EC50 10-fold lower in irradiated cells. These data further indicated that the clinical use of MRCK inhibition may have the greatest benefit when used in combination with RT. This radiation-specific reduction in pMRCK in response to MRCK inhibition was confirmed in an additional cell line (Fig. 6B). Importantly, subconfluent migration assays demonstrated that this reduction in MRCK activity translated into a reduction in radiation-driven migration across a panel of cell lines. Of note, no effect on viability was observed at the concentrations used in motility assays in two lines (1µM; Panc1, KPC). Interestingly, the patient derived TKCC04 line was sensitive to BDP9066 under both control and irradiated conditions indicating that in some conditions BDP9066 may have anti-tumorigenic properties in addition to inhibiting local invasion (Fig. 6C).

**Fig 6.**
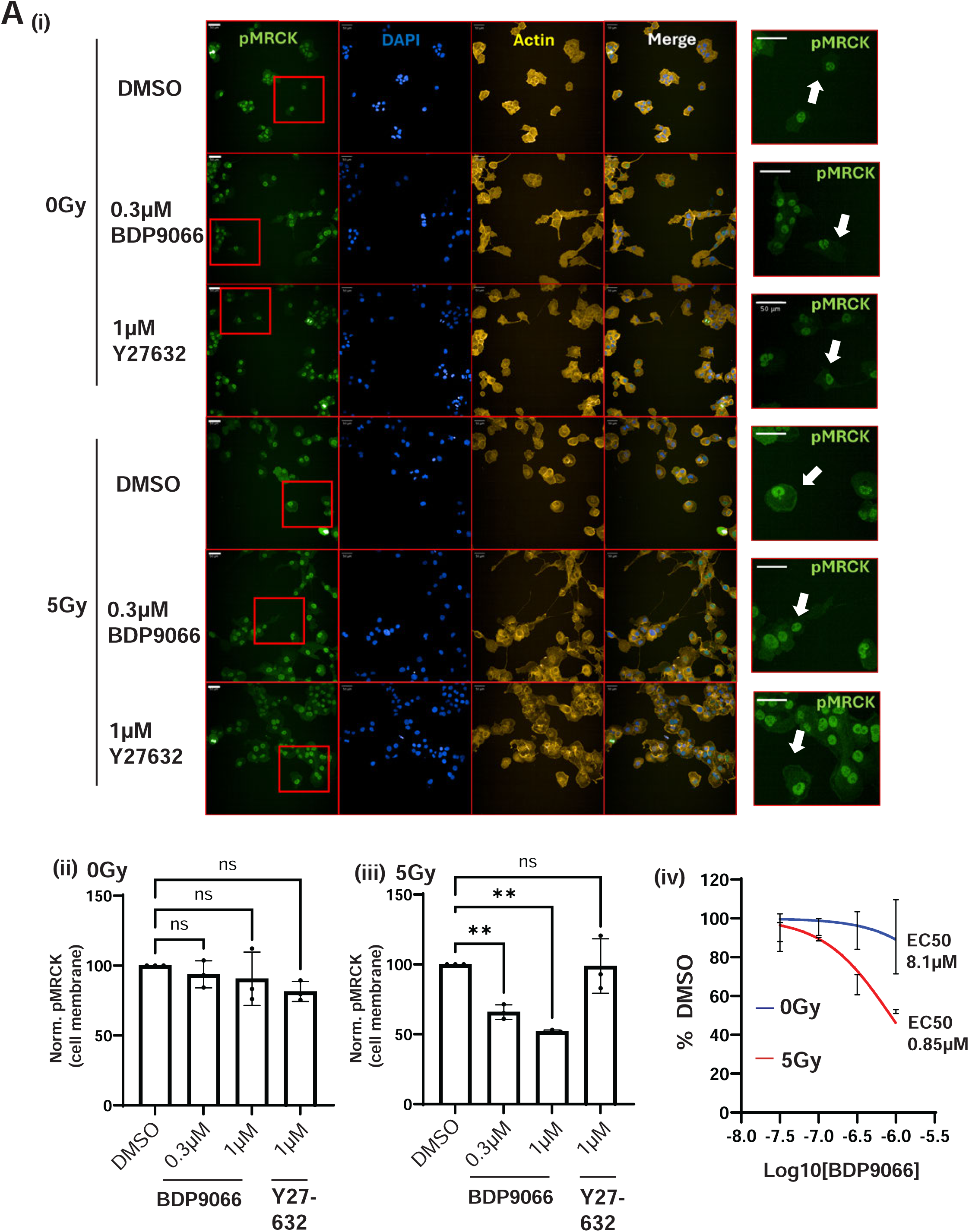
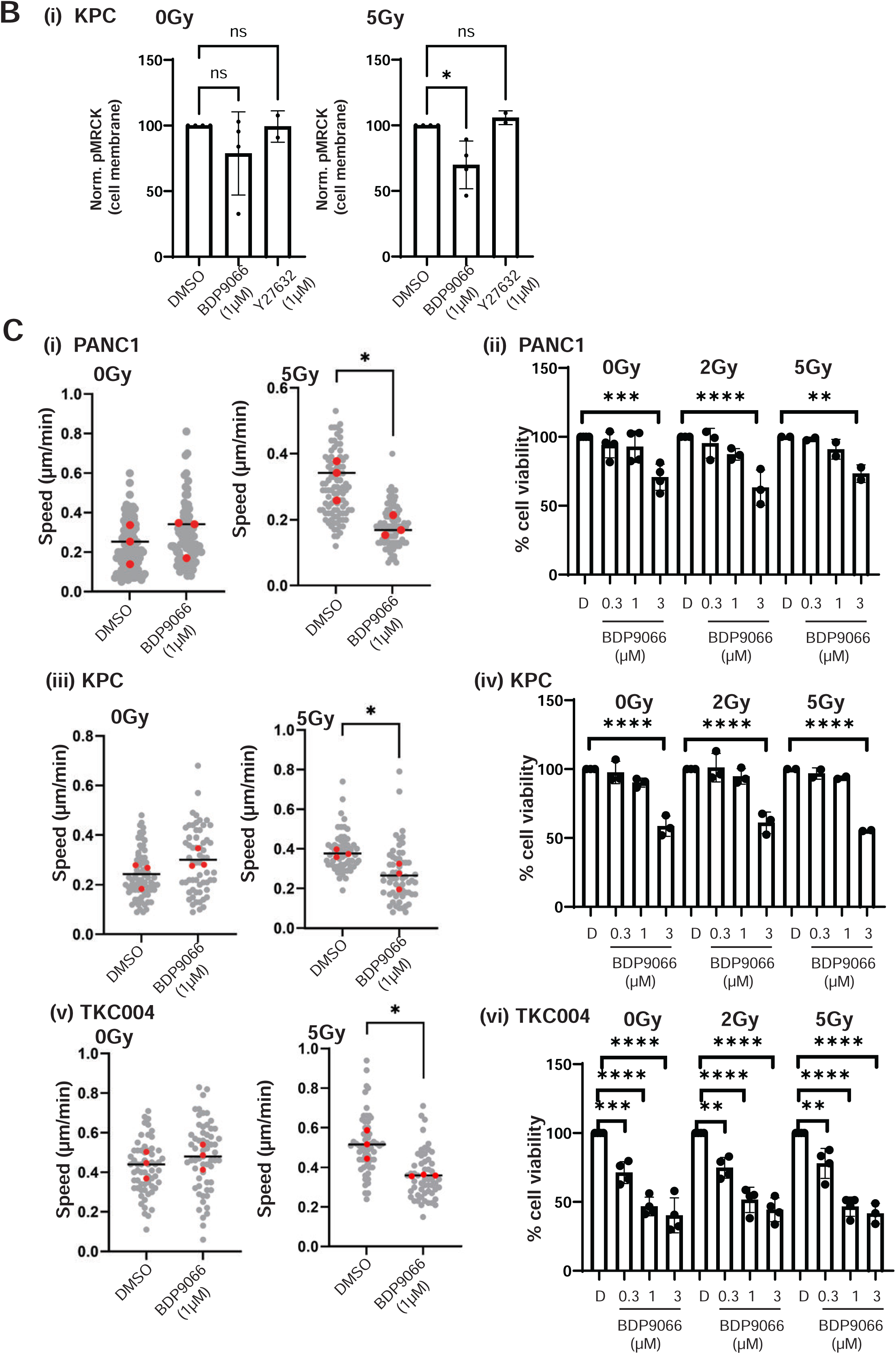
Pharmacological targeting of MRCK opposes radiation driven migration. **A)** (i) PANC1 cells treated with either 0 Gy or 5 Gy +/-DMSO, BDP9066 or Y27632 followed by high-throughput imaging. Blue, DAPI; yellow, actin; green, pMRCK; last panel, merge. Scale bar, 100 μm. White arrows indicate cell membrane. (ii)-(iii) Automated analysis of membrane associated pMRCK from (i). (iv) Dose response curve in the presence/absence of RT (5Gy). All data from 3 independent repeats. Statistical analysis: one-way ANOVA. ns, non-significant, **, *P* < 0.01. **B)** Automated analysis of membrane associated pMRCK in KPC (NBC12.2f) cells. Statistical analysis: one-way ANOVA. ns, non-significant, *, *P* < 0.05. Data from 3 independent experiments, error bars = standard deviation. **C)** Subconfluent migration and cell viability assays using (i)-(ii) PANC1, (iii)-(iv) KPC and (v)-(vi) TKC004 cells to measure the response to BDP9066 in the presence/absence of RT (5Gy). 0Gy and 5 Gy plotted separately due to experiment design. All data from 3 independent repeats. Statistical analysis: two-tailed, unpaired *t* test (motility) or one way anova (viability). *, *P <* 0.05 **, *P* < 0.01 ***, *P* < 0.005, ****, *P* < 0.001

### MRCK activity is upregulated at the invasive tumour margins and metastases post radiotherapy

To determine whether MRCK activity was upregulated post RT *in vivo*, we stained the sections from the KPC GEM model (Fig. 2) for pMRCK. While there was no increase in pMRCK across the tumor bulk (Fig. 7A(i)), pMRCK was significantly upregulated at the tumor margins compared to that at the tumor center (Fig. 7A (ii), (iii), (iv), (v)). Furthermore, this pattern of activation was concomitant with an increase in Zeb1 expression (a marker of epithelial-to-mesenchymal transition; Fig. 7B). These data indicate that RT may induce an MRCK-dependent, pro-invasive signature at the invasive borders of PDAC tumors. Importantly, this increase in pMRCK was maintained in metastases from the radiotherapy-treated cohort, indicating that MRCK signaling may be required to support progression from local invasion to distant metastases (Fig 7C). Finally, using *ex vivo* cultured tumor tissue from KPC GEM mice, we demonstrated that pMRCK levels, while not raised in response to RT across the tumour section (mirroring the *in vivo* response described above), is significantly reduced by BDP9066 when used in combination with RT, further highlighting the potential for therapeutic translation (Fig. 7D; Supplemental M&M).

**Fig 7.**
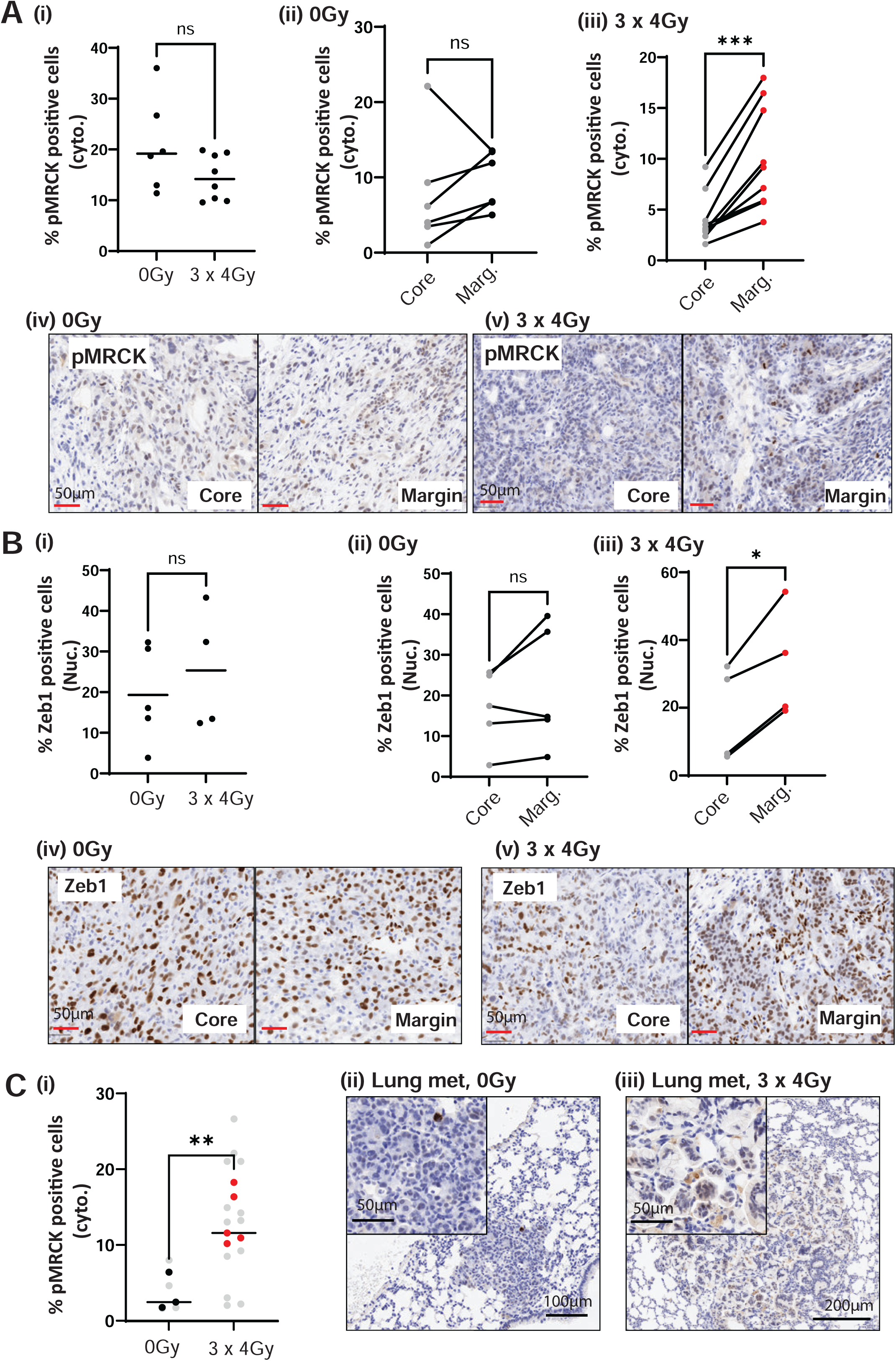
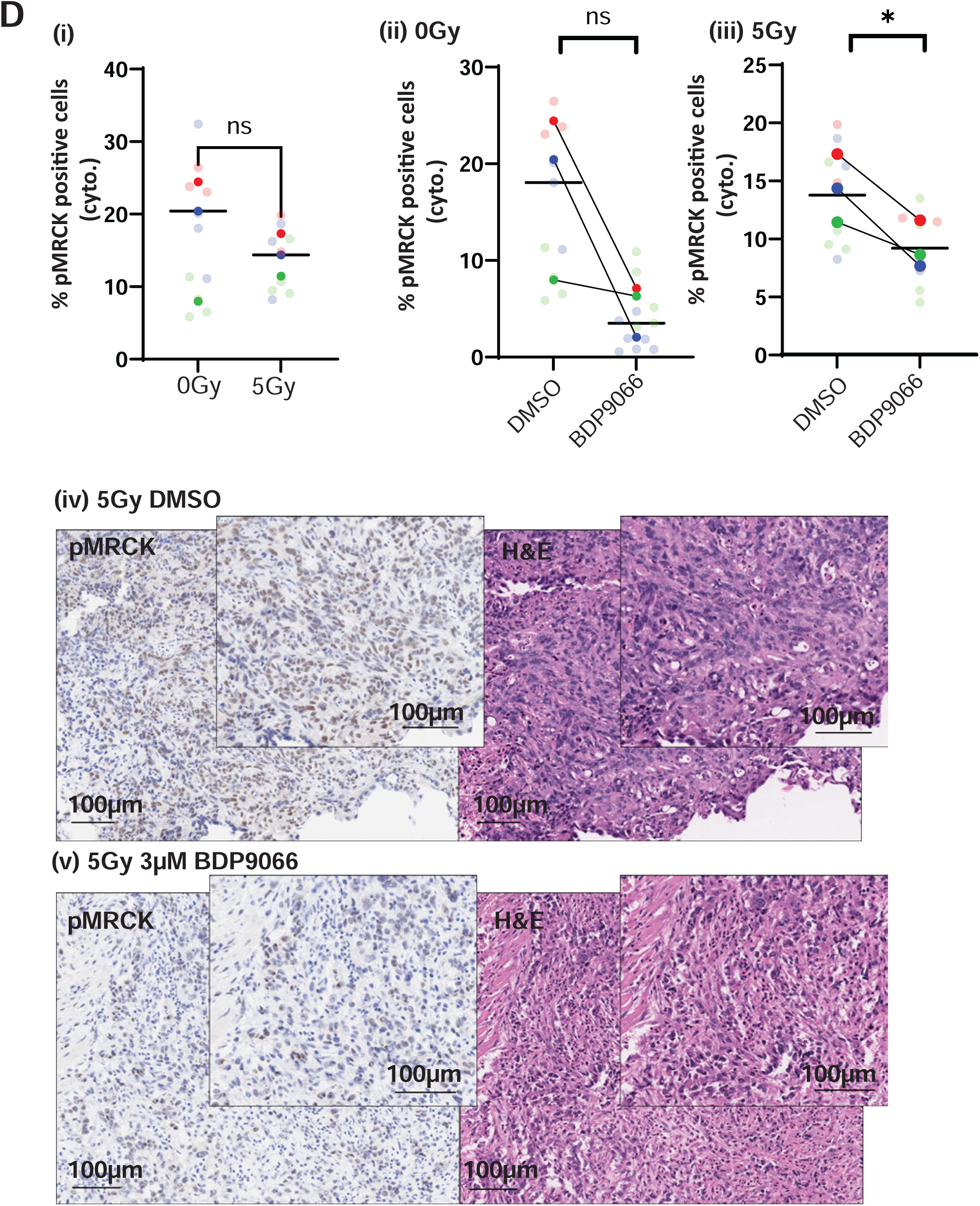
MRCK activity is upregulated at invasive tumour margins post radiotherapy *in vivo*. **A**) (i) Percentage of cytoplasmic pMRCK positive cells in KPC tumours treated with either 0 Gy or 3 x 4 Gy, n=6 and 8 mice, respectively. (ii, iii) Percentage of cytoplasmic pMRCK positive cells in the core v margin of KPC tumours treated with 0 Gy (ii) or 3 x 4Gy (iii). Means from individual tumours plotted. (iv, v) Example images from (ii)-(iii). Statistical analysis: two-tailed, unpaired *t* test. ns, non-significant, ***, *P <* 0.001. **B)** (i) Percentage of nuclear ZEB1 positive cells in KPC tumours treated with either 0 Gy or 3 x 4 Gy. (ii, iii) Percentage of nuclear Zeb1 positive cells in the core v margin of KPC tumours treated with 0 Gy (ii) or 3 x 4Gy (iii). (iv, v) Example images from (ii)-(iii). Means from individual mice plotted, n=5 0Gy, n=4 3×4Gy. Statistical analysis: two-tailed, unpaired *t* test. ns, non-significant, *, *P <* 0.05. **C)** (i) Percentage of cytoplasmic pMRCK positive cells in liver and lung metastases from KPC mice treated with 0 Gy or 3 x 4 Gy. Individual metastases from mice plotted in grey. Average per mouse in black/red. Statistical analysis: two-tailed, unpaired *t* test. *, *P <* 0.05. (ii, iii) Example images from (i). **D)** Percentage of pMRCK positive cells (cytoplasmic) in sections of 300µm slices from KPC tumours cultured *ex vivo* treated with 0 or 5Gy alone (i) or in combination with DMSO or BDP9066 (ii)-(ii), n=3 mice per group. (iv), (v) exemplar images. Statistical analysis: paired t test, ns, non-significant, *, *P <* 0.05. Means from individual experiments plotted as bold data points. Internal replicate data points plotted in corresponding transparent data points.

Together, our data indicate that RT drives increased metastasis in a clinically relevant model of PDAC, with organotropism towards the lung. This is driven, at least in part, by upregulation of MRCK signaling. Targeting MRCK with a small-molecule inhibitor may present a novel therapeutic avenue for containing spread of disease during radiotherapy.

## Discussion

Metastasis is a leading cause of morbidity and mortality in PDAC, with approximately 50% of patients having metastasis at diagnosis, increasing to 90% upon autopsy (36). Identifying a therapeutic window for the addition of anti-metastatic therapy has historically been a challenge for the cancer field. The clear induction of metastasis by targeted radiotherapy in a clinically relevant model of PDAC provides strong evidence for the first time that RT-driven metastasis may have an important clinical impact. Importantly, we observed increased metastasis within a short timeframe post-RT (two weeks) in animals that likely already had locally advanced disease at the time of treatment, mirroring the clinical situation. These data indicate a clear window of opportunity for the addition of anti-metastatic therapy concomitant with RT to control the dissemination rates and improve survival outcomes. Indeed, constraining metastatic spread may be predicted to become increasingly urgent as better methods for early, pre-metastatic detection of PDAC arise in the coming years. Although our data provide an important advance in understanding the effect of RT on metastasis, to fully understand the clinical relevance of our observations, the identification and analysis of outcome data from relevant patient cohorts should be performed in the future.

Surprisingly, our data indicate that not only does RT drive metastasis, but also a switch in organotropism towards the lungs. Although patients with isolated lung metastases have better survival outcomes, our data clearly indicate a potential therapeutic window whereby the addition of anti-metastatic therapy during RT may abrogate the metastatic spread of the disease and improve survival (37). The observed lung organotropism also provides strong evidence for an RT-specific switch in metastatic drivers. Indeed, our observation of increased pMRCK along with an increase in the mesenchymal marker Zeb1 at the tumor borders indicates a cell signaling switch towards MRCK-driven mesenchymal-like invasion during radiotherapy. Notably, single-cell RNA-seq data generated by our team from irradiated KPC tumors did not identify a clear EMT signature, which may suggest that the MRCK-driven response represents a partial EMT that is unique to the post RT scenario (data not shown). Unpicking this signaling response in the future has the potential to not only further our understanding of RT-driven metastasis but also of what drives lung-specific metastasis under physiological conditions in solid cancers. In addition, the surprising conservation of this response across cancers of starkly different origins (PDAC and glioblastoma) suggests that unpicking the surrounding signaling cascades will further our understanding of radiobiological responses (23).

Previous studies have demonstrated that RT drives more fibrotic stroma in solid cancers, including PDAC, which is largely driven by cancer-associated fibroblasts (CAFs) (9,38). This suggests that the switch to mesenchymal-like migration may be a necessary adaptation that allows PDAC cells to locally invade the post-RT landscape, resulting in increased metastatic spread. It is of interest that this switch even occurs *in vitro* in the absence of a stromal compartment, perhaps suggesting that the PDAC cells themselves have a key direct role in the deposition of fibrotic ECM post RT. Indeed, preliminary secretomic and proteomic data from our group indicated that PDAC cells have an altered ECM secretion profile post RT (data not shown).

Finally, we demonstrated the potential of utilizing this pro-metastatic switch in PDAC cells as a therapeutic window through the addition of a small molecule inhibitor of MRCK. Although BDP9066 is well tolerated *in vivo*, its short plasma half-life limits its translational potential (35). The development of a more stable iteration of the compound, or potentially the targeting of MRCK through alternative approaches, such as PROTAC, is key to moving this approach towards the clinic.

In summary, our data highlight the impact of RT on the metastatic landscape of PDAC and identify a novel, exploitable therapeutic approach to improve outcomes in patients with this cancer of current unmet need in the future.

## Data Availability

The authors confirm that the data supporting the findings of this study are available within the manuscript and its supplementary materials or are available from the authors on request.

## Author Contributions

**Conception and design:** KMM, MT, JPM, JLB

**Development of methodology:** KMM, MT, MO, JPM, JLB, LMC, RC, JM, AJC and LMcG.

**Acquisition of data (provided animals, acquired and managed patients, provided facilities, etc.):** KM, MT, LD, RC, DG, SL, JM AJC, and YS

**Analysis and interpretation of data (e.g., statistical analysis, biostatistics, computational analysis):** KM, MT, LD, RC, DG, YS, SL, RS, CM, NM and JM

**Writing, review, and/or revision of the manuscript:** JLB, KM, JM, MO, and MT.

No conflicts of interest declared.

## Acknowledgements

The histology was provided by the Histology facility of the CRUK-Scotland Institute. Imaging was provided by the Beatson Advanced Imaging Facility (RRID:SCR_023875) at the CRUK-Scotland Institute. *In vivo* support was provided by the Biological Service Unit of the CRUK Scotland Institute. Mass spectroscopy undertaken by the proteomics facility, CRUKSI and transcriptomics by the RNA sequencing facility, CRUKSI. Thank you to Prof. David Chang, Dr. Derek Grose and Dr. Aileen Dufton for lending their clinical insight into the review of this data.

