## Supplemental figs and legends for "Radiotherapy drives lung metastasis in PDAC via a RhoGTPase signaling shift towards MRCK-dependency"

**A**

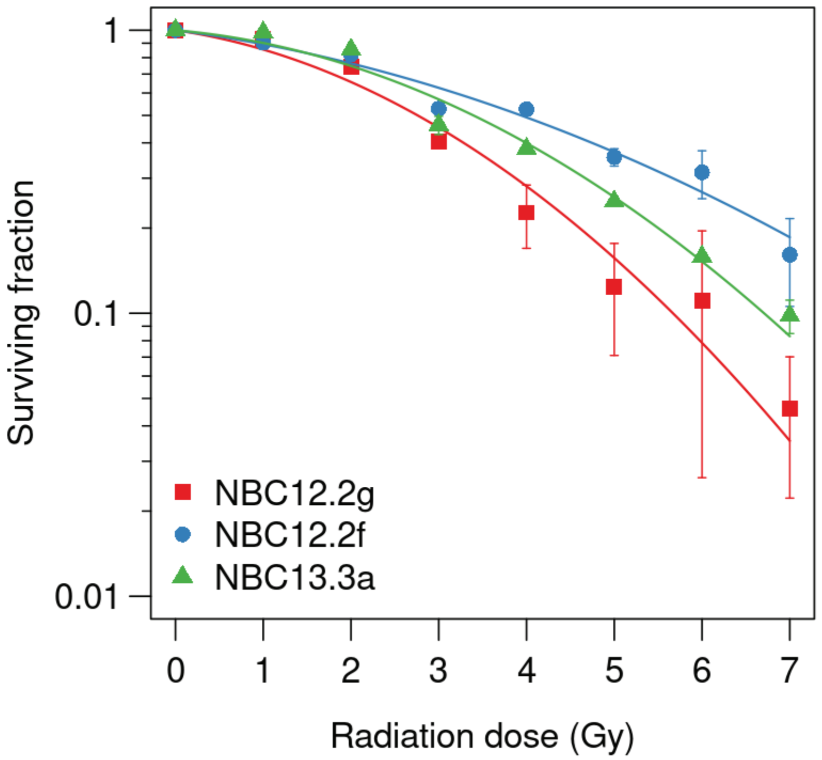
Surviving fraction
1
0.1
0.01
NBC12.2g
NBC12.2f
NBC13.3a
0
1
2
3
4
5
6
7
Radiation dose (Gy)

**Fig S1.**

**KPC cell lines exhibit varying degrees of radiosensitivity**

Clonogenic data for lines derived from KPC mice. Data from two independent repeats.

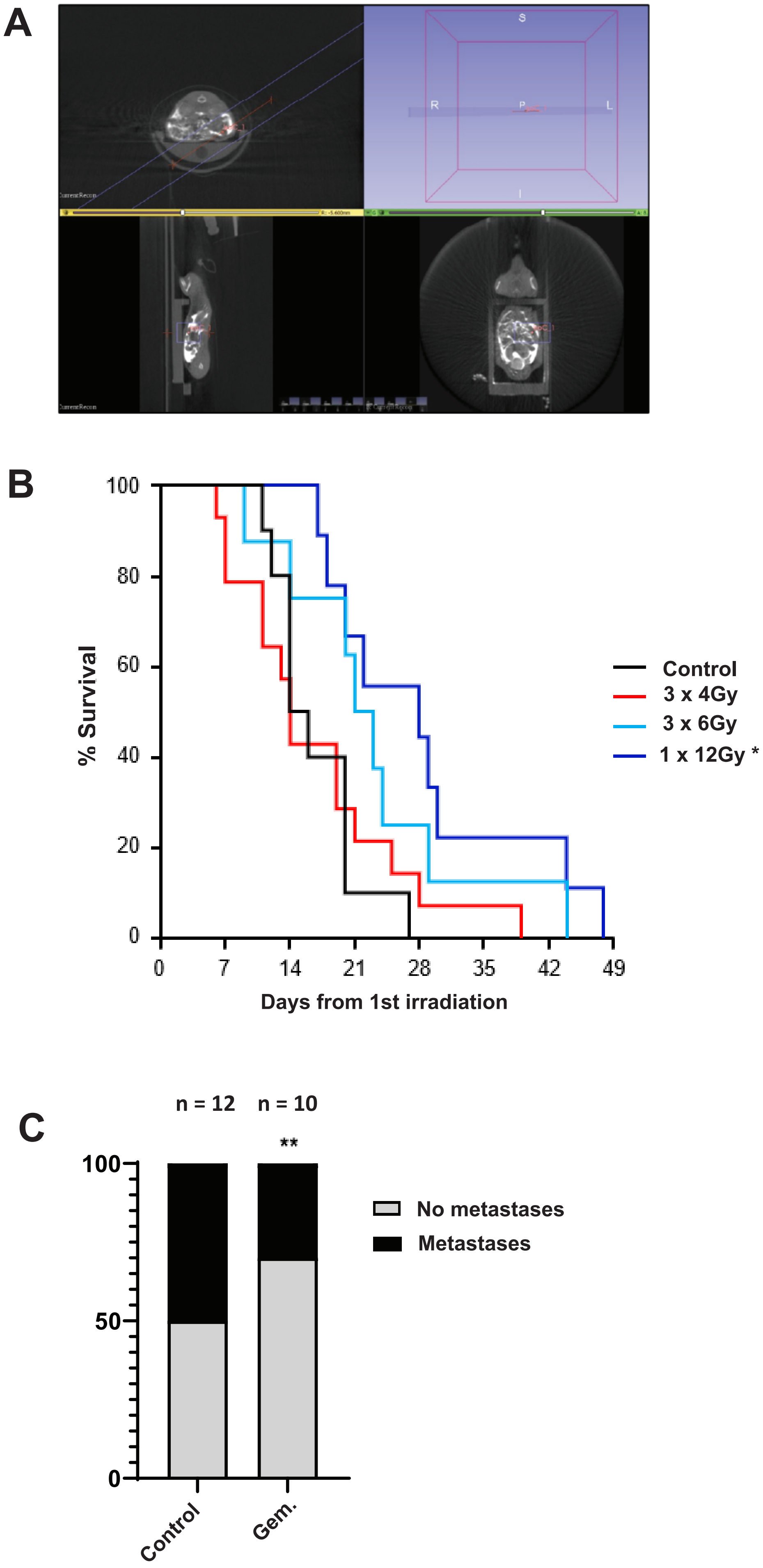
A
S
R
P
L
I
B
% Survival
100
80
60
40
20
0
0
7
14
21
28
35
42
49
Days from 1st irradiation
Control
3 x 4Gy
3 x 6Gy
1 x 12Gy *
C
n = 12
n = 10
**
100
50
0
Control
Gem.
No metastases
Metastases

**Fig 2.**

**Radiotherapy increases the metastatic burden of pancreatic cancer in a clinically relevant genetic mouse model. A)** Tumour bearing KPC mice were subjected to CT-guided targeted RT and sampled at clinical endpoint. Liver and lung sections were blinded and double scored for presence of metastases. **B) (i)** Serial sectioning protocol **(ii)**, **(iii)** example images of liver and lung metastases. **C)** i) H&E sections from KPC mice treated with 0Gy, 3x4Gy, 3x6Gy or 1x12Gy were scored for the presence of local invasion (example in (ii)). **D)** H&E liver, lung, and diaphragm sections from KPC mice treated with 0 or 3x4Gy RT were assessed for presence of metastases. **E)** Metastasis scoring across multiple RT protocols, as indicated. **F)** Breakdown of metastases present in liver **(i)** or lung **(ii)**. Number per cohort indicated on graphs. Statistical analysis: Chi-square test. ns, non-significant, *, $P < 0.05$, **, $P < 0.01$, ***, $P < 0.001$ ****, $P < 0.0001$.

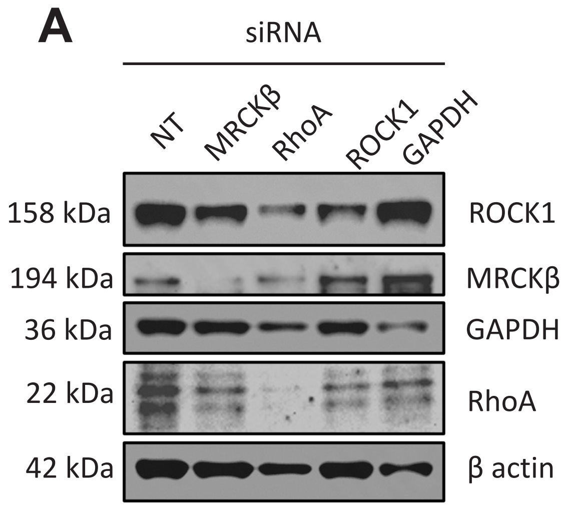
A
siRNA
NT
MRCKβ
RhoA
ROCK1
GAPDH
158 kDa
ROCK1
194 kDa
MRCKβ
36 kDa
GAPDH
22 kDa
RhoA
42 kDa
β actin

**Fig S3. Example western blot following siRNA treatment targeting ROCK1, MRCKβ, GAPDH and RhoA.** βactin used as loading control. NT= non targeting siRNA. Western blot run in parallel to screen to confirm knock down.
