## Supplemental Tables for "Radiotherapy drives lung metastasis in PDAC via a RhoGTPase signaling shift towards MRCK-dependency"

Table S1. Primary antibodies used for western blotting, immunofluorescences and immunohistochemistry

| Antibody | #Cat number | Dilution |
|---|---|---|
| MRCK | Prof Mike Olson, Ryeson University, Canada | 1/200 |
| pMRCK | Prof Mike Olson, Ryeson University, Canada | 1/500 |
| Zeb1 | Cell Signalling, #70512 | 1/500 |
| pMYPT1 | Millipore #ABS45 | 1/500 |
| b-Actin | Cell Signalling #3700S | 1/20 000 |
| GAPDH | Abcam #ab8245 | 1/2000 |
| ROCK1 | BD Transduction Laboratories, #611137 | 1/500 |
| CDC42BPA (MRCKα) | Abnova, #H00008476-M01 | 1/500 |
| CDC42BPA (MRCKα) | Abnova, H00009578-M03 | 1/500 |
| RhoA | Thermo Fisher Scientific #MA1-134 | 1/500 |

Table S2. siRNAs used in motility screen (Fig. 3)

| | Catalog Number, Dharmacon | Sequence |
|---|---|---|
| **RHOG** | L-008995-00 | CUACACAACUAACGCUUUC |
| **RHOG** | L-008995-00 | CCAGUCCGCCGUCCUAUGA |
| **RHOG** | L-008995-00 | GCUGUGCGCUACCUCGAAU |
| **RHOG** | L-008995-00 | GAGUAUGACCGCCUCCGUA |
| **CDC42BPB** | L-004075-00 | GCUUAGAGACCCAGAAUUG |
| **CDC42BPB** | L-004075-00 | GCUAUGAGAUCCAGAGAAC |
| **CDC42BPB** | L-004075-00 | GGAGGUGCAUGAUUCAGAA |
| **CDC42BPB** | L-004075-00 | GGUGACGGCCUCUCUCUUA |
| **RHOT1** | L-010365-01 | UGUGGAGUGUUCAGCGAAA |
| **RHOT1** | L-010365-01 | GCAAUUAGCAGAGGCGUUA |
| **RHOT1** | L-010365-01 | CCAGAGAGGGAGACACGAA |
| **RHOT1** | L-010365-01 | GCUUAAUCGUAGCUGCAAA |
| **RHOBTB2** | L-009252-00 | GACCGUCGCUUUGCUUAUG |
| **RHOBTB2** | L-009252-00 | CAUCCGAGCUGCACUCAUC |
| **RHOBTB2** | L-009252-00 | GGUAAGACCAGGCUCAUCU |
| **RHOBTB2** | L-009252-00 | CAGAAGAUCCUCUCACCUA |
| **RAC1** | L-003560-00 | GUGAUUUCAUAGCGAGUUU |
| **RAC1** | L-003560-00 | GUAGUUCUCAGAUGCGUAA |
| **RAC1** | L-003560-00 | AUGAAAGUGUCACGGGUAA |
| **RAC1** | L-003560-00 | GAACUGCUAUUUCCUCUAA |
| **RHOJ** | L-010367-00 | UCAAUUAUCUGAGGUUGUC |
| **RHOJ** | L-010367-00 | UCAUAGGGACCCAGAUUGA |
| **RHOJ** | L-010367-00 | UCAGAAAGGUCUCAAAGCG |
| **RHOJ** | L-010367-00 | AGAAACCUCUCACUUACGA |
| **RAC2** | L-007741-00 | UGACAACUAUUCAGCCAAU |
| **RAC2** | L-007741-00 | CCAAGGAGAUUGACUCGGU |

| | | |
|---|---|---|
| **RAC2** | L-007741-00 | CCAAGUGGUUCCCAGAAGU |
| **RAC2** | L-007741-00 | UGAAAACCGUGUUCGACGA |
| **RHOA** | L-003860-00 | CGACAGCCCUGAUAGUUUA |
| **RHOA** | L-003860-00 | GACCAAAGAUGGAGUGAGA |
| **RHOA** | L-003860-00 | GCAGAGAUAUGGCAAACAG |
| **RHOA** | L-003860-00 | GGAAUGAUGAGCACACAAG |
| **RHOT2** | L-008340-01 | GCGUGGAGUGUUCGGCCAA |
| **RHOT2** | L-008340-01 | CCUCAAGUUUGGAGCCGUU |
| **RHOT2** | L-008340-01 | GAGGUUGGGUUCCUGAUUA |
| **RHOT2** | L-008340-01 | AGGAGAUCCACAAGGCAAA |
| **RHOB** | L-008395-00 | GCAUCCAAGCCUACGACUA |
| **RHOB** | L-008395-00 | CAGAACGGCUGCAUCAACU |
| **RHOB** | L-008395-00 | CGACGAGCAUGUCCGCACA |
| **RHOB** | L-008395-00 | AAGCACUUCUGUCCCAAUG |
| **RND1** | L-008929-00 | GGAUCUCCCUACUACGAUA |
| **RND1** | L-008929-00 | GAGCUUAGUCUCUGGGAUA |
| **RND1** | L-008929-00 | AGACAGACCUGCGAACAGA |
| **RND1** | L-008929-00 | CAGAAGAGCCCUGUCCGAA |
| **CDC42** | L-005057-00 | CGGAAUAUGUACCGACUGU |
| **CDC42** | L-005057-00 | GCAGUCACAGUUAUGAUUG |
| **CDC42** | L-005057-00 | GAUGACCCCUCUACUAUUG |
| **CDC42** | L-005057-00 | CUGCAGGGCAAGAGGAUUA |
| **RND2** | L-009727-00 | GCGAUCCGCUCAGCUGUCA |
| **RND2** | L-009727-00 | CGACAUUAGCCGACCAGAA |
| **RND2** | L-009727-00 | ACGCCUAUCCCGGGAGUUA |
| **RND2** | L-009727-00 | ACACAAGGAUCGAGCCAAA |
| **ROCK1** | L-003536-00 | CUACAAGUGUUGCUAGUUU |
| **ROCK1** | L-003536-00 | UAGCAAUCGUAGAUACUUA |

| | | |
|---|---|---|
| **ROCK1** | L-003536-00 | CCAGGAAGGUAUAUGCUAU |
| **ROCK1** | L-003536-00 | GCCAAUGACUUACUUAGGA |
| **RHOD** | L-008940-00 | CGGCUCGGCUCCAUGACAA |
| **RHOD** | L-008940-00 | CCACGGUGUUUGAGCGGUA |
| **RHOD** | L-008940-00 | GCAAGAAGGUACCCAUCAU |
| **RHOD** | L-008940-00 | GAUUGGAGCCUGUGACCUA |
| **RHOC** | L-008555-00 | GAAAGAAGCUGGUGAUCGU |
| **RHOC** | L-008555-00 | GAACUAUAUUGCGGACAUU |
| **RHOC** | L-008555-00 | GGACAUGGCGAACCGGAUC |
| **RHOC** | L-008555-00 | CUACGUCCCUACUGUCUUU |
| **RHOH** | L-008804-00 | GAGUACAGCAGGUGUUUGA |
| **RHOH** | L-008804-00 | AACCAUAACUCAUUCCUGA |
| **RHOH** | L-008804-00 | GGACACAGCCGGCAAUGAC |
| **RHOH** | L-008804-00 | GAACUGCCGUCAACCAGGC |
| **RHOV** | L-006374-00 | GGACGAUGUCAACGUACUA |
| **RHOV** | L-006374-00 | GUAUUUGACUCGGCUAUUC |
| **RHOV** | L-006374-00 | GCUCAGCCUUGACGCAGAA |
| **RHOV** | L-006374-00 | GAAACUGAAUGCCAAAGGU |
| **RHOF** | L-008316-00 | CAUCGGUGUUCGAGAAGUA |
| **RHOF** | L-008316-00 | GCAAGACAGACCUGAGGAA |
| **RHOF** | L-008316-00 | UCCCUGAGGUCACGCAUUU |
| **RHOF** | L-008316-00 | UGAAGAAGGCGCAACGGCA |
| **CDC42BPA** | L-003814-00 | GCGCAAGACUCACCAGUUU |
| **CDC42BPA** | L-003814-00 | GACCAUACACUAUCAUUUA |
| **CDC42BPA** | L-003814-00 | GUUAGUAGCCCAACAGAUA |
| **CDC42BPA** | L-003814-00 | GUAACAGAAUCAAGUCAUU |
| **RHOQ** | L-009943-00 | CACCUAGAAUGUAAGUUAA |
| **RHOQ** | L-009943-00 | GCAGUUGGUCCCUAAGUGA |

| | | |
|---|---|---|
| **RHOQ** | L-009943-00 | GAGCGACAACUUAUAUACA |
| **RHOQ** | L-009943-00 | GUAAUAAGGUCAUAACUGC |
| **RHOBTB3** | L-020480-00 | ACAGAUGGCCGUCGAAUAU |
| **RHOBTB3** | L-020480-00 | CCGGAAAUGUCGUUGCUUA |
| **RHOBTB3** | L-020480-00 | GAAGAGUCCCACUGACAUU |
| **RHOBTB3** | L-020480-00 | CCAUGAACCUUGAUAUAGU |
| **RAC3** | L-008836-00 | AAACUGACGUCUUUCUGAU |
| **RAC3** | L-008836-00 | ACAAGAAGCUGGCACCCAU |
| **RAC3** | L-008836-00 | GAAGACAUGCUUGCUGAUC |
| **RAC3** | L-008836-00 | CGUGAUGGUGGACGGGAAA |
| **RND3** | L-007794-00 | CUACAGUGUUUGAGAAUUA |
| **RND3** | L-007794-00 | UAGUAGAGCUCUCCAAUCA |
| **RND3** | L-007794-00 | CAGCAAAUCUAUCAUGGAU |
| **RND3** | L-007794-00 | GCGGACAGAUGUUAGUACA |
| **RHOBTB1** | L-009389-00 | GAACUUGGCUUACCAUACU |
| **RHOBTB1** | L-009389-00 | GGACGUGACAUUUAAAUUG |
| **RHOBTB1** | L-009389-00 | GAACACCCGUUAUCCUUGU |
| **RHOBTB1** | L-009389-00 | GACAGACGCUUUGCAUAUG |
| **RHOU** | L-009882-00 | CAUCGUCGCUGGCAUUCAA |
| **RHOU** | L-009882-00 | AAGCAGGACUCCAGAUAAA |
| **RHOU** | L-009882-00 | GUACUGCUGUUUCGUAUGA |
| **RHOU** | L-009882-00 | GAACGUCAGUGAGAAAUGG |
| **ON-TARGETplus Non-targeting Control** | D-001810-10 | UGGUUUACAUGUCGACUAA |
| **ON-TARGETplus Non-targeting Control** | D-001810-10 | UGGUUUACAUGUUGUGUGA |

| | | |
|---|---|---|
| **ON-TARGETplus Non-targeting Control** | D-001810-10 | UGGUUUACAUGUUUUCUGA |
| **ON-TARGETplus Non-targeting Control** | D-001810-10 | UGGUUUACAUGUUUUCCUA |

Table S3. Summary of literature examining the effects of radiotherapy on PDAC cell motility and invasion

| Article | Cell line | In vitro/ex vivo/in vivo | Dose | Migration | Invasion |
|---|---|---|---|---|---|
| **Fujita et al., 2011** | Mia PaCa-2, PANC-1 | In vitro | 2 Gy/4 Gy | Unchanged PANC-1, enhanced Mia PaCa-2 ($p<0.05$) | Enhanced invasion in both cell lines ($P<0.05$) |
| **Ohuchida et al., 2004** | Suit-2 | In vitro | 5 Gy | | Unchanged |
| **Qian et al., 2002** | PANC-1, Suit-2, Hs766T | In vitro | 3 Gy/5Gy /10 Gy | Reduced in PANC-1 and Suit-2, unchanged Hs766T | Enhanced PANC-1, Suit-2 ($P<0.05$), unchanged Hs766T |
| **Fujita et al., 2012** | PANC-1 | In vitro | Carbon-ion – 0, 0.5, 1, 2 or 4 Gy | | Enhanced PANC-1 ($P<0.05$), reduced Mia PaCa-2, BxPC-3 and AsPC-1 |
| **Yao et al., 2011** | PANC-1, Mia PaCa-2, BxPC-3 | In vitro | 1-6 Gy | | Enhanced PANC-1, Mia PaCa-2 and BxPC-3 |
