## Supplemental Materials and Methods for "Radiotherapy drives lung metastasis in PDAC via a RhoGTPase signaling shift towards MRCK-dependency"

**Proteomics**

Proteins were reduced with 10mM DTT and subsequently alkylated in the dark with 55 mM Iodoacetamide, both reactions were carried out at room temperature for one hour. Alkylated proteins were precipitated adding 4 volumes of Acetone at -20°C overnight. Washed pellets were reconstituted in 50 µl of HEPES 200 mM and digested first with Endoproteinase Lys-C (ratio 1:33 enzyme:lysate) for one hour, followed by an overnight trypsin digestion (ratio 1:33 enzyme:lysate).

The digested peptides from each experiment were differentially labelled using TMT16-plex reagent (Thermo Scientific). The label reagents 127N, 128N, 129N, 130N, 131N, 132N, 133N, 134N were used to label secretome samples, whereas the label reagents 126, 127C, 128C, 129C, 130C, 131C, 132C, 133C were used to label proteome samples. Each sample was labelled with 0.1 mg of TMT reagent dissolved in 50 µl of 100% anhydrous acetonitrile. The reaction was carried out at room temperature for 2 hours. Fully labelled samples were separately mixed in equal amount and desalted using 50 mg Sep Pak C18 reverse phase solid-phase extraction cartridges (Waters). 63

TMT-labelled peptides from were fractionated using high pH reverse phase chromatography on a C18 column (150 × 2.1 mm i.d. - Kinetex EVO (5 µm, 100 Å)) using an HPLC chromatography system (Agilent, LC 1260 Infinity II, Agilent). A two-step gradient of 30 and 60 minutes were applied to separate TMT labelled peptides from secretome and proteome respectively. Column eluates were collected in 21 fractions directly analysed for proteome sample or pooled into 10 fractions for secretome samples.

Peptides resulting from all samples were separated by nanoscale C18 reverse-phase liquid chromatography using an EASY-nLC II 1200 (Thermo Scientific) coupled to an Orbitrap Q-Exactive HF (Thermo Scientific) for proteome samples, or to an Orbitrap Fusion Lumos (Thermo Scientific) for secretome samples. Elution was carried out at a flow rate of 300 nl/min using a binary gradient with buffer A (2% acetonitrile) and B (80% acetonitrile), both containing 0.1% formic acid. Samples were loaded with 6 µl of buffer A into a 50 cm fused silica emitter (New Objective) packed in-house with ReproSil-Pur C18-AQ, 1.9 µm resin (Dr Maisch GmbH). For both systems the packed emitter was kept at 50 °C by means of a column oven (Sonation) integrated into the nanoelectrospray ion source (Thermo Scientific), and the Xcalibur software (Thermo Scientific) was used for data acquisition.

Peptides were eluted using different gradients optimised for three sets of fractions: 1–7, 8–15, and 16–21 (Cao et al., 2020). Each fraction was acquired for a duration of 190 minutes. A full scan over mass range of 350–1400 m/z was acquired at 120,000 resolution at 200 m/z. Higher energy collisional dissociation fragmentation was performed on the most intense ions during 3 sec cycle time. Ions were selected within an isolation window of 0.8 m/z, and peptide fragments were analysed in the Orbitrap at 15,000 resolution.

The Mass-Spec Raw data were processed with MaxQuant software (Cox and Mann, 2008) version 1.6.14.0 (Proteome) or 2.3.0.0 (Secretome) and searched with Andromeda search engine (Cox et al., 2011). First and main searches were performed with precursor mass tolerances of 20 ppm and 4.5 ppm, respectively, and MS/MS tolerance of 20 ppm. The minimum peptide length was set to six amino acids and specificity for trypsin cleavage was required, allowing up to two missed cleavage sites. The peptide, protein, and site false discovery rate (FDR) was set to 1 %. Modification by iodoacetamide on Cysteine residues (Carbamidomethylation) were specified as fixed, whereas Methionine oxidation and N-terminal acetylation modifications were specified as variable. MaxQuant was set to quantify on “Reporter ion MS2”, and interference between TMT channels were corrected by MaxQuant using the correction factors provided by the manufacturer. The “Filter by PIF” option was activated and a “Reporter ion tolerance” of 0.003 Da was used.

From both experiments, the proteinGroups.txt file from MaxQuant output was used for protein quantitation analysis using Perseus software version 1.6.15.0 (Cox and Mann, 2008). The “Reverse”, “Potential Contaminants” and “Only identified by site” protein, as specified in MaxQuant, were removed, as well as protein groups identified with no unique peptides. Only proteins robustly quantified in all replicates in at least one group, were allowed in the list of quantified proteins. Significantly enriched proteins were selected using a permutation-based Student’s t-test or ANOVA with FDR set at 5%.

*“The raw files and the MaxQuant search results files have been deposited as complete submission to the ProteomeXchange Consortium [1] via the PRIDE partner repository [2] with the dataset identifier PXD068500.”*

*[1] = Deutsch EW, Bandeira N, Perez-Riverol Y, Sharma V, Carver J, Mendoza L, Kundu DJ, Wang S, Bandla C, Kamatchinathan S, Hewapathirana S, Pullman B, Wertz J, Sun Z, Kawano S, Okuda S, Watanabe Y, MacLean B, MacCoss M, Zhu Y, Ishihama Y, Vizcaíno JA (2023). The ProteomeXchange Consortium at 10 years: 2023 update. Nucleic Acids Res, 51(D1):D1539-D1548 (Pubmed ID: 36370099).*

*[2] = Perez-Riverol Y, Bai J, Bandla C, Hewapathirana S, García-Seisdedos D, Kamatchinathan S, Kundu D, Prakash A, Frericks-Zipper A, Eisenacher M, Walzer M, Wang S, Brazma A, Vizcaíno JA (2022). The PRIDE database resources in 2022: A Hub for*

*mass spectrometry-based proteomics evidences. Nucleic Acids Res 50(D1):D543-D552 (PubMed ID: 34723319).*

### ***Ex vivo*** **tumour culture**

Tumour samples were taken from Pdx1-Cre; LSL-KrasG12D/+; LSL-Trp53R172H/+ (KPC) mice. The KPC mice were bred at the CRUK Scotland Institute and genotyped by Transnetyx (Cordoba, TN, USA).

Tumour samples were resected and transported to the tissue culture room within 15 minutes, samples were preserved in PBS containing 1 × antibiotic/antimycotic solution (on ice) during transportation. Resected samples were cut into 300μm section using 5100mz Vibratome (Campden Instruments). Sections were cultured on Millicell® cell culture inserts (Millipore, Ref. PICM03050) in 6-well plates at 37 °C, 5% $CO_2$ in a humidified environment. Dulbecco's Modified Eagle's Medium- high Glucose (500 ml, Merck, Ref. D5796), supplied with 10% FBS (Gibco, Thermo Fisher Scientific, Ref. A5256701), 0.01 mg/mL hydrocortisone (Merck, Ref. H0396), 0.01 mg/mL insulin (Merck, Ref. I9278-5ML) and 1 × Antibiotic Antimycotic Solution (100×, Merck, Ref. A5955-20ML) was used to supply nutrients, medium was replaced every day. Irradiation and/or drug treatment were applied on Day 0 after sections were allowed to equilibrate for 2 hours. Sections were fixed and submitted for histology on Day 3. 0 or 5Gy irradiation was delivered using the RS225 radiation cabinet.

### **RNAseq and bioinformatic analysis**

Analysis of RNA-Seq expression data M45_2301_irradiation_PDAC

Quality checks and trimming on the raw RNA-Seq data files were done using FastQC version 0.11.9(1), FastP version 0.20.1(2) and FastQ Screen version 0.15.1 (3). RNA-Seq paired-end reads were aligned to the GRCm39.108 version of the mouse genome and annotation (4), using HiSat2 version 2.2.1 (5) and sorted using Samtools version 1.14 (6).

Aligned genes were identified using Feature Counts from the SubRead package version 2.0.1 (7).

Expression levels were determined and statistically analysed using the R environment version 4.2.3 (8) and utilizing packages from the Bioconductor data analysis suite (9). Differential gene expression was analysed based on the negative binomial distribution using the DESeq2 package version 1.36.0 (10), batch corrected using ComBat from the sva package version 3.44.0 (11) and adaptive shrinkage using Apeglm version 1.18 (12).

Salmon (13) version 1.9.0 was used for the Quantification of transcripts. Fishpond version 2.4.1 (14) was used to determine Differential Transcript Expression and Differential Transcript Usage.

Identification of enriched biological functions was achieved using g:Profiler version 0.2.1 (15).

Computational analysis was documented at each stage using MultiQC version 1.14 (16), Jupyter Notebooks(17) and R Notebooks(18).

Robin Shaw

https://orcid.org/**0000-0001-5503-5233**
